# Glucose-Responsive CD20+ Cytotoxic T Cells: A Novel Pro-inflammatory Mediator in the Immunopathogenesis of Type 2 Diabetes

**DOI:** 10.64898/2026.02.12.705577

**Authors:** Özgür Albayrak, Adnan Batman, Selen Ünlü, Nazan Akkaya, Sarah Shamim Khan, Selin Çakmak Demir, Seçil Özışık, Havva Sezer, Zeynep Yağmur Yılmaz, Berk Mızrak, Oğuzhan Deyneli, Dilek Yazıcı, Atay Vural

## Abstract

**Objective:** Although T-cell-mediated inflammation is a hallmark of Type 2 Diabetes (T2D), the contribution of CD20-expressing T cells—a highly potent subset recently implicated in various autoimmune conditions—to T2D pathogenesis remains unknown. This study aimed to characterize the frequency, functional profile, and glucose-responsiveness of CD20+ cytotoxic T lymphocytes (CTLs) in patients with T2D.

**Research Design and Methods:** Peripheral blood mononuclear cells (PBMCs) from 25 treatment-naïve T2D patients and 20 age-matched healthy controls (HC) were analyzed using multicolor flow cytometry. We assessed the frequency of CD3+CD8+CD20+ cells and their production of cytotoxic molecules (Granzyme B, Perforin, Granzyme K) and pro-inflammatory cytokines (IFN-γ, TNF-α, GM-CSF). To further establish the link with hyperglycemia, HC-derived CTLs were exposed to increasing glucose concentrations (100–450 mg/dL) in vitro. Single-cell RNA sequencing (scRNA-seq) data from a public T2D dataset was utilized to validate the molecular signature of MS4A1 (CD20)+ CTLs.

**Results:** The frequency of circulating CD20+ CTLs was significantly elevated in T2D patients compared to HCs (p<0.0001) and demonstrated a strong positive correlation with HbA1c, fasting glucose, and triglyceride levels. Notably, CD20+ CTLs from T2D patients exhibited a “hyperfunctional” phenotype, characterized by significantly higher degranulation (CD107a), elevated expression of Granzymes B/K and Perforin, and increased production of IFN-γ and TNF-α compared to HCs (p<0.01 for all). In contrast, no such differences were observed in the CD20-CTL compartment. In vitro experiments revealed that escalating glucose levels directly enhanced the proliferation and cytotoxic potential of CD20+ CTLs, suggesting a nutrient-sensing mechanism. scRNA-seq analysis further confirmed the distinct pro-inflammatory and effector transcriptional profile of MS4A1+ T cells in T2D.

**Conclusions:** Our findings identify CD20+ CTLs as a novel, glucose-sensitive, and hyperfunctional immune subset in T2D. The strong correlation between these cells and clinical metabolic parameters suggests that CD20+ CTLs may act as a critical link between chronic hyperglycemia and systemic inflammation, representing a potential new therapeutic target for immunomodulation in T2D.

## INTRODUCTION

According to the World Health Organization (WHO), diabetes is one of the four non-communicable diseases with the highest mortality rates (1). Type 2 diabetes (T2D), is the most common form of diabetes, occurring when body cells resist and inadequately utilize the insulin produced (2). T2D can cause macrovascular complications such as coronary artery disease, peripheral artery disease, cerebrovascular disease, and microvascular complications such as retinopathy, nephropathy, and neuropathy (3).

The pathophysiology of T2D involves several interrelated mechanisms, including activation of the hexosamine biosynthetic pathway, accumulation of advanced glycation end products (AGEs), and the stimulation of signaling cascades via the protein kinase C (PKC) pathway, all of which are promoted by oxidative stress in a hyperglycemic environment. (4) There is also a crucial role of abnormal immune cell activation and due to this, subsequent inflammation plays a crucial role in the progression of T2D. (5–7)

Numerous studies have shown impaired T cell functions in individuals with T2D. Peripheral blood mononuclear cells (PBMCs) isolated from diabetic patients with obesity were observed to produce lower amounts of IL-2, IL-6, and TNF-α upon stimulation with phytohemagglutinin (PHA) (8). Martinez et al. demonstrated a decrease in pathogen-specific memory Th17 responses and the number of CD4+ T cells in patients with diabetes in response to stimulation with Streptococcus pneumoniae (9). Kumar et al. also investigated the functions of CD8+ T cells and NK cells in the whole blood of T2DM patients infected with Mycobacterium tuberculosis. Compared to controls, the patients exhibited a reduction in cytokine production (IFN-γ, IL-2, IL-17A/F, and TNF-α) and the expression of cytotoxic molecules (perforin, granzyme B, and CD107a) by CD8+ and NK cells (10, 11).

Although the expression of the CD20 molecule was long thought to be restricted to B cells, subsequent studies have shown that this molecule is also expressed, albeit at low levels, by T cells (referred to as CD3+CD20dim or CD3+CD20+). Interestingly, T cells expressing CD20 exhibit a more activated phenotype compared to CD3+CD20-cells, are elevated in effector memory T cell compartment and have a higher CD8/CD4 ratio Notably, these cells have been found to proliferate faster and earlier than B cells following rituximab treatment. (12–14)

The role of CD20+ T cells in endocrinological diseases were only studied in two studies, so far. A study by Stramazzo et.al found that CD8+CD20+ T lymphocytes are significantly elevated in Hashimoto’s thyroiditis—especially in hypothyroid and polyautoimmune cases—and their increased frequency correlates with worse thyroid function, suggesting these proinflammatory cells actively contribute to autoimmune thyroid tissue damage and disease progression (15). Study by Pinho et.al. also showed that, elevated CD20+ T cell levels and expression in peripheral blood and adipose tissue are linked to obesity-related adiposity, inflammation, insulin resistance, and cardiometabolic risk and Type 2 Diabetes, suggesting their potential as markers of complications, particularly in pre-BMS individuals. (16) However, there are no studies regarding the direct role of CD20+ CTLs cells in T2DM.

In this study, we comprehensively characterized the frequency, effector function, and metabolic responsiveness of CD20+ CTLs in treatment-naïve T2D patients compared with healthy controls. We further investigated how increasing glucose concentrations modulate proliferation and cytotoxic profile of CD20+ CTLs of healthy controls. Furthermore, by utilizing the single-cell RNA-sequencing dataset from the study by Gu et al., titled “Single-cell analysis of human PBMCs in healthy and type 2 diabetes populations: Dysregulated immune networks in type 2 diabetes unveiled through single-cell profiling,” we conducted differential gene expression (DGE) analysis between MS4A1□ and MS4A1□ cytotoxic T cells. This allowed us to characterize the molecular differences of CD20□ cytotoxic T cells in T2D relative to their CD20□ counterparts and to further investigate the association of these cells with T2D (17). Our findings uncover a hyperfunctional, glucose-responsive CD20+ CTL subset that may serve as a novel link between hyperglycemia, inflammation, and immune-mediated tissue injury in T2D.

## RESEARCH DESIGN and METHODS

### Study Population and PBMC Isolation

Twenty-five newly diagnosed, treatment-naïve adults with Type 2 diabetes (T2D) and twenty age- and sex-matched healthy controls were enrolled. All T2D participants were non-smokers, free of acute or chronic infections, autoimmune disorders, malignancy, or corticosteroid use. Healthy controls had no metabolic or systemic inflammatory conditions. Inclusion criteria comprised: age 18–65 years, absence of glucose-lowering therapy, and no prior immunomodulatory medication. Exclusion criteria included pregnancy, acute illness, hepatic or renal impairment, and known hematologic disease. All participants provided written informed consent. The study protocol was approved by the Koç University Institutional Review Board with the decision number 2023.31 0.IRB 1.106. Peripheral blood was collected in EDTA tubes (K3EDTA (BD Biosciences, USA) from T2D patients and healthy volunteers. PBMCs were isolated using density gradient centrifugation method using Lymphoprep (d=1.077g/ml, Axis-Shield, Norway). After isolation, PBMCs were cryopreserved in liquid nitrogen until the day of the experiments. On the day of the experiments, cryopreserved PBMCs were thawed at 37^0^C water bath and taken into complete medium (RPMI 1640 + 10% FBS + 1% Penicillin-Streptomycin, GIBCO, USA). All the centrifugation steps were performed at 500 xg for 5 minutes unless reported otherwise. All antibodies were purchased from Biolegend (Biolegend, USA). On the day of the experiments, cryopreserved PBMCs were thawed at 37^0^C water bath and taken into complete medium (RPMI 1640 + 10% FBS + 1% Penicillin-Streptomycin, GIBCO, USA). All the centrifugation steps were performed at 500 xg for 5 minutes unless reported otherwise.

### Immunophenotyping and Functional Studies of CD20+ Cytotoxic T Cells

PBMCs were seeded into 96 well round bottom plates at a density of 1 x 10^6^ cells/100 ul RPMI medium with or without 1X Cell Activation Cocktail (81nM PMA + 1.3µM Ionomycin without Brefeldin A, Biolegend, USA) containing mouse anti-human CD107a Brilliant Violet 421 antibody (Clone H4A3) and stimulated for 1 hour at 37^0^C 5% CO_2_ incubator. After 1 hour, Brefeldin A (Biolegend, USA) was added into the wells, and an additional stimulation was carried out for 3 hours. After stimulation, plates were centrifuged, supernatants were discarded, and cells were resuspended and incubated with PBS containing Zombie NIR fixable viability dye (Biolegend, USA) for 10 minutes at RT in dark. After incubation, plates were centrifuged, supernatants were discarded, and cells were resuspended with 100ul staining buffer containing CD3 FITC (Clone HIT3a), CD8 Alexa Fluor 700 (Clone SK1), CD20 APC (Clone 2H7), CD19 APC-Cy7 (Clone HIB19), and CD14 APC-Cy7 (Clone HCD14) antibodies and incubated at RT in dark for 20 minutes. All antibodies were purchased from Biolegend (Biolegend, USA). After incubation, plates were centrifuged, and cells were fixed with Fixation Buffer and permeabilized by Permeabilization Wash Buffer (Biolegend, USA) according to the manufacturer’s instructions. After fixation/permeabilization, cells were resuspended with permeabilization buffer containing mouse anti-human Granzyme B PE-Cyanine5 (Clone QA16A02), Perforin PE (Clone B-D48), Granzyme K PE-Cyanine7 (Clone GM26E7) (Panel 1) and IFN-γ PE (Clone B27), TNF-α PE-Cyanine7 (Clone MAB11), rat anti-human GM-CSF PerCP-Cyanine 5.5 (Clone BVD2-21C11) (Panel 2) and incubated for 20 minutes at RT in dark. After incubation, plates were washed with permeabilization buffer and staining buffer respectively, resuspended with staining buffer, transferred into 15×75mm tubes, and analyzed with Beckman Coulter Cytoflex SRT flow cytometer (Beckman Coulter, USA). Flow cytometric analysis was done with FlowJo v 10.10.0 (BD Biosciences, USA). Gating Strategy for the detection of CD20+ T cells are given in Supplementary Figure 1 (**Figure S1**). Detailed information about the antibodies (fluorochrome, clone, brand) used in the experiments are given in Supplementary Table 1.

### Proliferation of CD20+ Cytotoxic T Cells

To compare the proliferation of CD20+ CTLs from T2D patients and healthy controls, PBMCs were labeled with Tag-It Violet proliferation tracking dye according to the manufacturer’s protocol. Subsequently, cells were incubated with PBS containing Zombie NIR fixable viability dye (Biolegend, USA) for 10 minutes at RT in dark, washed with staining buffer, and resuspended with 100ul staining buffer containing CD3 FITC (Clone HIT3a), CD8 Alexa Fluor 700 (Clone SK1), CD20 PE (Clone 2H7), CD19 APC-Cy7 (Clone HIB19), and CD14 APC-Cy7 (Clone HCD14) antibodies and incubated at RT in dark for 20 minutes. After incubation, cells were washed with staining solution, resuspended in RPMI medium, and CD19-CD14-CD4-CD3+CD8+CD20+ and CD19-CD14-CD4-CD3+CD8+CD20-T cells were sorted using Cytoflex SRT cell sorter (Beckman Coulter, USA) with purity setting and a speed of ≤2000 events/second and experiment proceeded with samples exhibiting >95% purity (**Figure S2)**. 1×10^5^ cells were seeded into 96 well round bottom plates and cultured in 100 µL of media containing 5 µg/mL PHA, 10 ng/mL IL-2, and 10 ng/mL IL-15 for 96 hours at 37°C with 5% CO2. After stimulation, plates were centrifuged, supernatants were discarded, cells were resuspended with staining buffer, transferred into 15×75mm tubes, and analyzed with Beckman Coulter Cytoflex SRT flow cytometer (Beckman Coulter, USA). Flow cytometric analysis was done with FlowJo v10.10.0 (BD Biosciences, USA).

### Proliferation and Functions of CD20+ Cytotoxic T Cells in Increasing Glucose Concentrations

To investigate the impact of elevated glucose concentrations on the proliferation and function of CD20+ cytotoxic T cells, PBMCs and sorted CD8+CD20+ and CD8+CD20-cells from healthy controls were exposed to increasing glucose concentrations for 96 hours. For this purpose, Glucose-Free RPMI medium (Gibco, USA) was supplemented with D-glucose (Sigma-Aldrich, Germany) at concentrations of 100 mg/dL, 200 mg/dL, 300 mg/dL, 350 mg/dL, 400 mg/dL and 450 mg/dl.

For proliferation assays, a fraction of PBMCs was labeled with Tag-It Violet (Biolegend, USA) according to manufacturer’s instructions. Subsequently, cells were incubated with PBS containing Zombie NIR fixable viability dye (Biolegend, USA) for 10 minutes at RT in dark, washed with staining buffer, and resuspended with 100ul staining buffer containing CD3 FITC (Clone HIT3a), CD8 Alexa Fluor 700 (Clone SK1), CD20 PE (Clone 2H7), CD19 APC-Cy7 (Clone HIB19), and CD14 APC-Cy7 (Clone HCD14) antibodies and incubated at RT in dark for 20 minutes. After incubation, cells were washed with staining solution, resuspended in RPMI medium, and CD19-CD14-CD4-CD3+CD8+CD20+ T and CD19-CD14-CD4-CD3+CD8+CD20-T cells were sorted using Cytoflex SRT cell sorter (Beckman Coulter, USA) with purity setting and a speed of ≤2000 events/second and experiment was continued with samples with >95% purity (**Figure S2)**. 5×10^4^ CD8+CD20+ and CD8+CD20-cells were cultured at each glucose concentration in 100 µL of media containing DynaBeads Human T Cell Activation kit (Thermo-Scientific, USA), 10 ng/mL IL-2, and 10 ng/mL IL-15 for 96 hours at 37°C with 5% CO2. Remaining fractions of healthy control PBMCs were seeded at a cell density of 2.5×10^5^ cells/100ul for each glucose concentration containing DynaBeads Human T Cell Activation kit (Thermo-Scientific, USA), 10 ng/mL IL-2, and 10 ng/mL IL-15 for 96 hours at 37°C with 5% CO_2_. After the incubation, unsorted PBMCs were stained with the fixable viability dye and antibodies mentioned above to determine the changes in the CD8+CD20+ cell population and CD20 expression.

After stimulation, plates were centrifuged, and supernatants from sorted CD8+CD20+ and CD8+CD20-cells were collected for determining the concentrations of cytotoxic molecules, specifically Granzyme B, Granzyme A, and Perforin, along with pro-inflammatory cytokines IFN-γ and TNF-α secreted by CD20+ cytotoxic T cells, using LegendPlex Human CD8/NK Kit (Biolegend, USA) according to the manufacturer’s protocol. Samples were run with Attune NxT flow cytometer (Thermo Scientific, USA) and analyzed with LegendPlex Qognit analysis software (Biolegend, USA). In order to measure Granzyme K concentration Human Granzyme K ELISA Kit (Abcam, USA) was used according to manufacturer’s instructions and concentrations were determined using MultiSkan GO Multiplate Spectrophotometer (Thermo Scientific, USA)

Cell pellets were resuspended with staining buffer, transferred into 15×75mm tubes, and analyzed with Beckman Coulter Cytoflex SRT flow cytometer (Beckman Coulter, USA). Flow cytometric analysis was done with FlowJo v10.10.0 (BD Biosciences, USA).

### scRNA-seq data analysis

Peripheral blood single cell RNA-sequencing data from 37 individuals with type 2 diabetes (17) were obtained from Gene Expression Omnibus with GSE268210 accession number and processed in Scanpy (18). Doublets were identified with Scrublet (19) and removed. Cells with high mitochondrial transcript fraction (percent mitochondrial genes ≥ 20%) and with excessive complexity (> 5,000 detected genes) were excluded. Counts were library-size normalized to 10,000 per cell and log-transformed [log (x + 1)]. Gene expression was standardized to zero mean and unit variance (values clipped at ±10), followed by principal component analysis (PCA). Batch correction was performed with Harmony (20) and the neighborhood graph was computed using the first 30 Harmony-corrected principal components (PCs). Clustering of the whole dataset was done with the Leiden algorithm (resolution = 0.5) to identify major cell types (21). Clusters were annotated with CellTypist (model: “Immune_All_High”), then manually curated (22, 23). T cells were subset and re-analyzed following the same pipeline as above to separate CD4 and CD8 compartments; contaminating cells in each compartment were removed based on canonical markers and CellTypist labels. MS4A1□ T cells were defined as those with nonzero MS4A1 expression (counts > 0). Expression of classical B-cell markers (BANK1, CD79A, CD79B, JCHAIN, CD19, CD40) and T-cell markers (CD3D/E/G, CD8A/B, CD4) were inspected in CD4 and CD8 compartments to confirm true T cell identity of MS4A1□ subsets (**Figure S6**). After isolation, CD8 T cell compartment was reprocessed following the same normalization, PCA, batch correction and neighborhood graph computation as above and cells were clustered with the Leiden algorithm at resolution 0.3. Differential expression between MS4A1□ and MS4A1□ CD8 T cells was performed using Scanpy’s “sc.tl.rank_genes_groups” function, with Wilcoxon rank-sum statistics and Benjamini–Hochberg correction. Over-representation analysis (ORA) with Enrichr was performed via gseapy package in Python on the significantly differentially expressed genes from the CD8 MS4A1 comparison (upregulated and downregulated genes were analyzed separately, FDR < 0.05) (24, 25). The following libraries were queried: GO Biological Process (2025), GO Cellular Component (2025), GO Molecular Function (2025), Reactome (2024), KEGG (2021 Human), MSigDB Hallmark (2020), BioCarta (2016), HDSigDB Human (2021), and GeneSigDB. Enrichment P values (Enrichr/Fisher’s exact) were Benjamini–Hochberg adjusted per library.

### tSNE Analysis of CD20+ CTLs within Memory, Granzyme K and Granzyme B Subsets

PBMCs of 20 T2D patients and 20 healthy controls were seeded into 96 well round bottom plates at a density of 1 x 10^6^ cells/100 ul PBS containing Zombie NIR fixable viability dye (Biolegend, USA) and incubated for 10 minutes at RT in dark. After incubation, plates were centrifuged, supernatants were discarded, and cells were resuspended with 100ul staining buffer containing CD3 FITC (Clone HIT3a), CD8 Alexa Fluor 700 (Clone SK1), CD20 APC (Clone 2H7), CCR7 PE (Clone G043H7), CD45RO BV650 (Clone UCHL1), CD19 APC-Cy7 (Clone HIB19), and CD14 APC-Cy7 (Clone HCD14) antibodies and incubated at RT in dark for 20 minutes. After incubation, plates were centrifuged, and cells were fixed with Fixation Buffer and permeabilized by Permeabilization Wash Buffer (Biolegend, USA) according to the manufacturer’s instructions. After fixation/permeabilization, cells were resuspended with permeabilization buffer containing mouse anti-human Granzyme B PE-Cyanine5 (Clone QA16A02) and Granzyme K PE-Cyanine7 (Clone GM26E7) antibodies and incubated for 20 minutes at RT in dark. After incubation, plates were washed with permeabilization buffer and staining buffer respectively, resuspended with staining buffer, transferred into 15×75mm tubes, and analyzed with Beckman Coulter Cytoflex SRT flow cytometer (Beckman Coulter, USA). Flow cytometric analysis was done with FlowJo v 10.10.0 (BD Biosciences, USA).

To visualize the distribution and frequencies of CD20+ cytotoxic T cells within Granzyme K and Granzyme B subsets, FCS files from T2D patients and healthy controls were concatenated within their respective groups using the ‘FCS Merge’ function of the Floreada.io online flow cytometry analysis software (https://floreada.io). Subsequently, GzmK+, GzmB+, GzmK+GzmB+ (double-positive), and double-negative populations were gated within the memory subsets of CD8+ compartment and CD20+ cells residing within these specific subsets were then identified and analyzed with the t-SNE function within FlowJo v10.10.0 (BD Biosciences, USA).

### Statistical Analysis

All statistical analyses were conducted using GraphPad Prism (version 10.2.3). Data distributions were assessed using the Shapiro–Wilk and Kolmogorov–Smirnov tests. As most immunophenotypic and cytokine variables showed non-Gaussian characteristics, non-parametric tests were applied throughout unless otherwise indicated. Group comparisons between T2D patients and healthy controls were performed using the Mann–Whitney U test. Correlations between CD20□ T-cell subsets and clinical parameters (HbA1c, fasting glucose, LDL cholesterol, triglycerides, BMI) were analyzed using Spearman’s rank correlation. Two-Way ANOVA test with Bonferroni’s correction was used to compare the functional parameters, and proliferative capacities of CD20+ and CD20-CTLs and distribution of CD20+ CTLs in Granzyme K+ and Granzyme B+ subsets within memory compartments between T2D patients and healthy controls. RM-One-Way ANOVA with Bonferroni’s correction was used to determine the changes in the functionality and proliferative capacities of CD20+ and CD20-CTLs in different glucose concentrations.

Receiver operating characteristic (ROC) analyses were used to test the discriminatory ability of CD20□ CTL frequencies and for identifying T2D; area under the curve (AUC) was calculated. For single-cell RNA-sequencing analyses, differential gene expression between MS4A1□ and MS4A1□ CD8□ T-cell subsets was assessed using the Wilcoxon rank-sum test with Benjamini–Hochberg correction. Only genes with adjusted p<0.05 were considered significant. All analyses were two-tailed, and a p-value < 0.05 was considered statistically significant.

## RESULTS

### Clinical and Biochemical Profiles of Study Participants

Enrolled patients were included into three groups based on their HbA1c levels: mild (between 5.8% and 6.5%, n=5), moderate (between 6.6% and 7.9%, n=5), high (between 8% and 9.9%, n=5), severe (between 10% and 11%, n=6) and critical (≥11%, n=4). Statistical analyses were done to compare clinical parameters between T2D patients and healthy controls and all HbA1c, FBG, LDL, triglyceride and BMI levels were found to be significantly elevated in T2D patients compared to healthy controls (**Figure S3A**). As a result, we expanded our clinical focus to include LDL, triglyceride, and BMI levels in addition to HbA1c and FBG in the study. The age, gender, and clinical information of the patients and controls participating in the study are presented in Table 1. The clinical and demographic characteristics of all T2D patients and the healthy control group are also presented in Supplementary

**Table 1.** Baseline Demographic, Clinical, and Biochemical Characteristics of Participants. Values are presented as mean ± SEM or median [IQR] where appropriate. Comparisons between T2D patients and healthy controls were performed using the Mann–Whitney U test. T2D: Type 2 Diabetes, HbA1c: Glycosylated Hemoglobin, FBG: Fasting Blood Glucose, LDL: Low-Density Lipoprotein, HDL: High Density Lipoprotein, BMI: Body Mass Index, AST: Aspartate Aminotransferase, ALT: Alanine Aminotransferase, CRP: C-Reactive Protein, eGFR: Estimated Glomerular Filtration Rate

|  | *Healthy Control* | *T2D* | *P* |
| --- | --- | --- | --- |
| *N* | 25 | 25 |  |
| *Age* | 50.08 ± 1.53 | 52.56 ± 1.26 | 0.2246 |
| *Sex* |  |  |  |
| *Male* | 17 | 16 |  |
| *Female* | 8 | 9 |  |
| *HbA1c %* | 5.47 ± 0.027 | 8.88 ± 0.48 | <0.0001 |
| *FBG mg/dl* | 91.88 ± 1.26 | 181.7 ± 13.3 | <0.0001 |
| *LDL mg/dl* | 107.6 ± 2.67 | 181.2 ± 18.2 | 0.0004 |
| *Triglyceride mg/dl* | 104.4 ± 4.66 | 188.2 ± 23.83 | 0.0004 |
| *HDL mg/dl* | 56.88 ± 5.6 | 49.76 ± 3.2 | 0.1591 |
| *BMI kg/m<sup>2</sup>* | 22.59 ± 1.78 | 29.41 ± 3.84 | <0.0001 |
| *AST u/L* | 22.76 ± 1.77 | 21.40 ± 2.01 | 0.1153 |
| *ALT u/L* | 21.16 ± 1.68 | 22.76 ± 1.42 | 0.2728 |
| *ALP u/L* | 58.76 ± 2.68 | 62.72 ± 3.67 | 0.2363 |
| *CRP mg/L* | 2.052 ± 0.47 | 5.092 ± 1.9 | 0.1042 |
| *Serum Creatinine mg/dl* | 0.81 ± 0.04 | 0.84 ± 0.08 | 0.8309 |
| *eGFR ml/min/1.73m<sup>2</sup>* | 106.9 ± 1.77 | 102.2 ± 1.97 | 0.1244 |

### Increased Frequency of Circulating CD20□ T Cells in Patients With Type 2 Diabetes

The frequency of circulating CD20□ T cells was significantly higher in patients with T2D compared with healthy controls. Among total CD3□ T cells, CD20□ cell percentages and CD20 mean fluorescence intensity (MFI) were both elevated in T2D among total CD3+ T cells (CD20% median IQR 9.81 [9.0-12.69], CD20 MFI median IQR 7962 [7477-8837]) as well as among CD4+ helper T cells (THLs) (CD20% median IQR 7.62 [6.075-9.075], CD20 MFI IQR 7964 [7287-8765]), and CD8+ cytotoxic T cells (CTLs) (CD20% median IQR 17.20 [11.20-19.85] CD20 MFI median IQR 8189 [7624-9212]) (**Figure 1A and 1B**). The discriminative value of this measure for T2D was very prominent, as revealed by the high area under curve values obtained by ROC analysis (**Figure 1C**).

**Figure 1.**
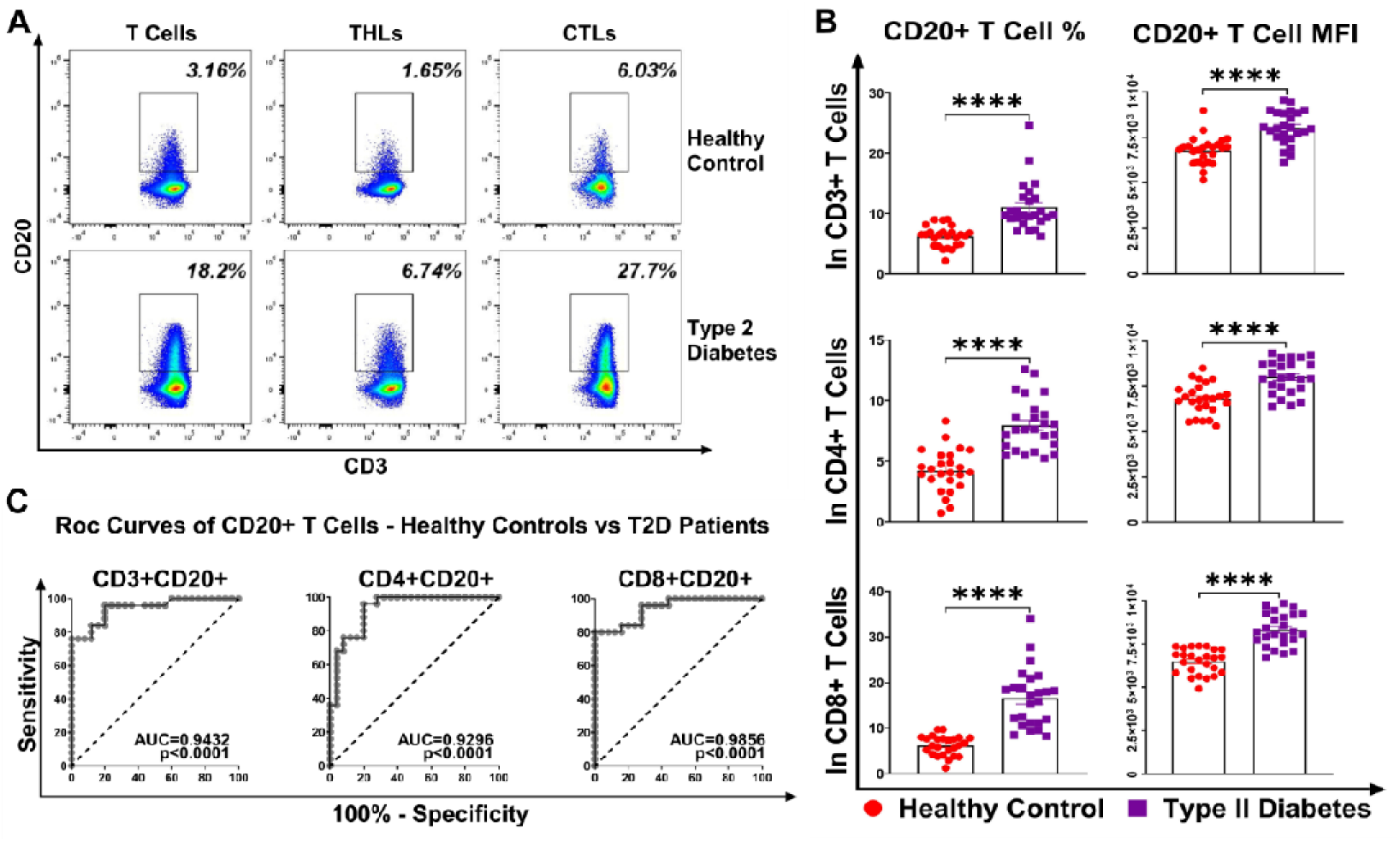
Comparative Analysis of Circulating CD20□ T Cells in Patients With Type 2 Diabetes and Healthy Controls. Statistical analysis was performed using the Mann-Whitney U test. **A and B)** Analysis of CD3+, CD3+CD4+, and CD3+CD8+ cells showed that the percentages of CD20+ T cells and CD20 MFI were significantly higher in T2D patients compared to healthy controls. **C)** ROC curve analysis was used to calculate AUC values for CD3+CD20+, CD4+CD20+, and CD8+CD20+ cells in healthy controls and T2D patients. *\*p<0.05, **p<0.01, ***p<0.001, ****p<0.*0001.

### Association of CD20□ T-Cell Frequencies With Metabolic Parameters

We performed correlation analysis to evaluate the relationship between CD20+ T cell percentages and the clinical profile of T2D. This analysis was restricted to HbA1c, FBG, LDL, Triglyceride, and BMI levels, as these were exclusively the parameters that differed significantly between the patient and control groups. We did not observe a correlation between the percentages of total CD20+ T cells and CD20+ THLs cells with HbA1c, FBG, LDL cholesterol, and triglyceride levels whereas, CD20+ CTLs demonstrated a positive correlation with these measured parameters. Despite obesity being one of the most significant risk factors for T2D pathogenesis and the presence of a significantly elevated BMI in our patient cohort compared to healthy controls, it is noteworthy that we did not detect any correlation between the BMI values of T2D patients and the percentages of CD3+CD20+, CD4+CD20+, or CD8+CD20+ T cells. (**Figure 2)**. Furthermore, we extended our correlation analysis to include clinical parameters that did not show statistically significant differences between T2D patients and healthy controls, such as HDL, ALP, AST, ALT, CRP, serum creatinine, and eGFR. (**Figure S4**).

**Figure 2.**
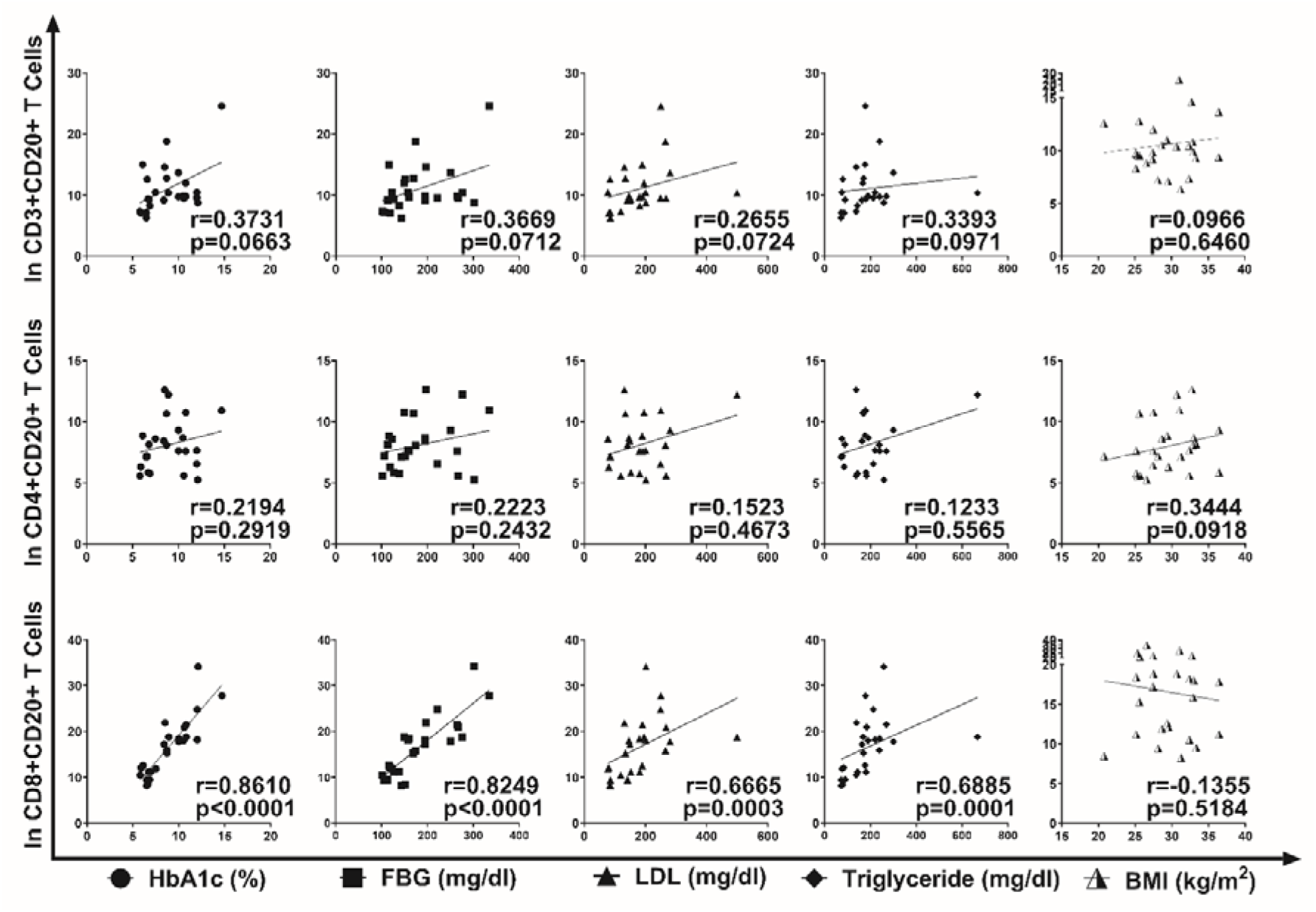
Correlation analyses between peripheral blood CD3+CD20+, CD3+CD4+CD20+, and CD3+CD8+CD20+ cell percentages and Metabolic Parameters in T2D. Spearman’s non-parametric correlation test was used for the analyses. Spearman’s r values were calculated, and p<0.05 was considered statistically significant.

### CD20□ Cytotoxic T Cells in T2D Patients Display Enhanced Degranulation and Cytotoxicity

Since the subgroup with the highest CD20+ population among T cells, yielded the most significant results in ROC curve analyses, and demonstrated a positive correlation with patients’ HbA1c, FBG, LDL, and triglyceride levels, we established our focus on the comparison of CD20+ CTL functions between T2D patients and healthy controls as the next phase of our study. To achieve this, we stimulated PBMCs of 20 T2D patients and 20 healthy controls with PMA/I for 4 hours.

Post stimulation analyses revealed that frequency of degranulated (Surface CD107a+) CD20+ CTL population was significantly elevated in patients with T2D compared to healthy controls (p=0.0001). Additionally, in the analysis of mean fluorescence intensity, we observed that the expression of CD107a was greater in the CD20+ CTLs of T2D patients (p<0.0001) We then separately compared CD20+ and CD20-CTLs in both healthy controls and T2D patients. Our analyses revealed that CD20+ CTLs exhibited greater degranulation compared to CD20-cells in both healthy controls and T2D patients (CD107a+% p<0.0001, CD107a MFI p<0.0001). However, when comparing CD20-CTLs, we observed no significant difference between T2D patients and healthy controls. (**Figure 3A and 3B**).

**Figure 3.**
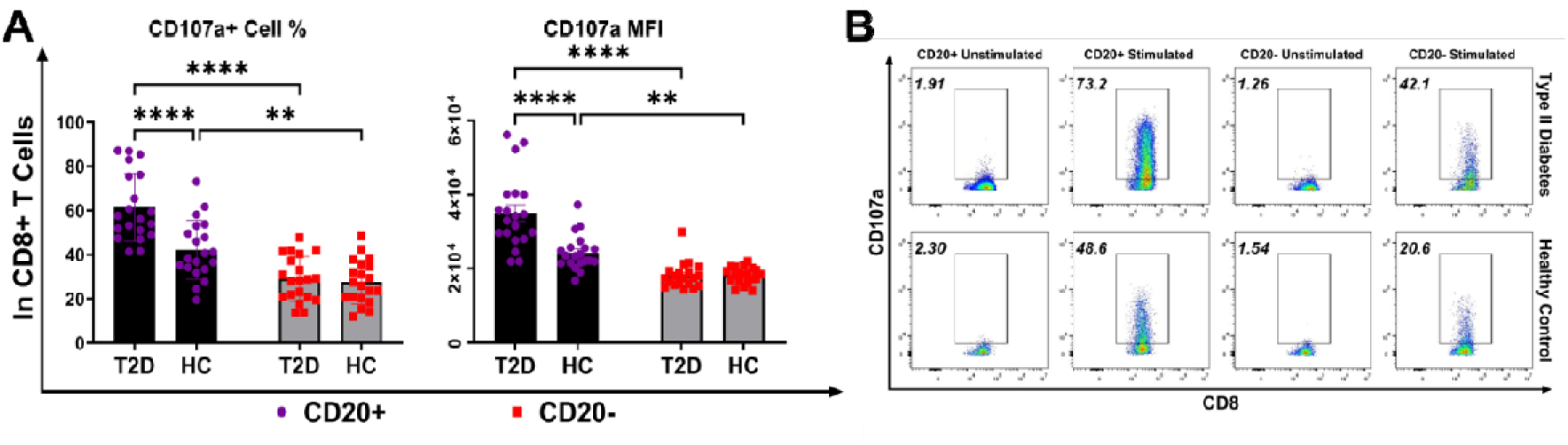
**A)** Enhanced Degranulation of CD20□ CD8L T Cells in Type 2 Diabetes based on the percentages of surface CD107a+ cells and CD107a MFI values. Statistical analysis was performed using Two-Way ANOVA test with Bonferroni correction. CD8+CD20+ T cells were found to exhibit higher cytotoxic characteristics compared to CD20-cells, regardless of disease status. No significant differences in the cytotoxic potential of CD8+CD20-cells were observed between T2D patients and healthy controls. However, CD8+CD20+ T cells in T2D patients demonstrated significantly higher cytotoxicity compared to those in healthy controls. **B)** Representative flow cytometric analysis of CD20+ and CD20-CTLs in T2D patients and healthy controls *\*p<0.05, **p<0.01, ***p<0.001, ****p<0.*0001.

Subsequently, in addition to degranulation, we examined the cytotoxic molecules Granzyme B and Perforin within CD20+ and CD20-CTLs in both T2D patients and healthy controls. In healthy controls, the Granzyme B+ cell population remained unchanged between CD20+ and CD20-cells whereas Perforin+ population were found to be elevated in CD20-compartment (p<0.05). Notably, a significant increase in Granzyme BL cell frequency was observed in CD20□ CTLs of T2D patients compared to controls (p < 0.0001), while Perforin^+^ percentages remained unchanged between groups (p<0.0001). When analyzing the MFI values of Granzyme B and Perforin, we observed higher expression levels in CD20+ CTLs compared to CD20-cells in both healthy controls (p<0.05) and T2D patients (p<0.0001) for granzyme B. No change was observed for Perforin MFI between CD20+ and CD20-CTLs of healthy controls whereas elevated Perforin expression was observed in CD20+ CTLs of T2D patients (p<0.0001). Additionally, a comparison of CD20+ cells between T2D patients and healthy controls revealed significantly higher Granzyme B and Perforin expression in patients. However, no differences were detected between CD20-cells of patients and controls. Similar to the cell percentages, we did not detect any differences in MFI values of Granzyme B and Perforin between the CD20-CTLs of T2D patients and healthy controls. Upon analyzing the Granzyme K molecule, we observed the following results: Both healthy controls and T2D patients exhibited a significantly higher percentage of Granzyme K+ cells (p<0.0001) and elevated Granzyme K expression (p<0.0001) in CD20+ CTLs compared to CD20-cells. Furthermore, when comparing CD20+ CTLs between T2D patients and healthy controls, we found that the Granzyme K+ cell population and Granzyme K expression were markedly and significantly elevated in the CD20+ CTLs of T2D patients. Once again, we found no differences in either Granzyme K+ cell percentages or Granzyme K MFI values between the CD20-cells of T2D patients and healthy controls (**Figure 4A and 4C**). In the final stage, we examined and compared the cells producing the pro-inflammatory cytokines IFN-γ, TNF-α, and GM-CSF, as well as the expression levels of these cytokines, within the CD20+ and CD20-cytotoxic T cells of T2D patients and healthy controls. Our analyses revealed that the percentages of IFN-γ+, TNF-α+, and GM-CSF+ cells, as well as their expression levels, were significantly higher in the CD20+ CTLs compared to CD20-cells in both healthy controls and T2D patients (Cell % p<0.001, MFI p<0.05). Additionally, when comparing T2D patients to healthy controls, we found that the CD20+ cells of T2D patients had significantly higher percentages of IFN-γ+, TNF-α+, and GM-CSF+ cells and elevated expression levels of these cytokines (Cell % p<0.01, MFI p<0.05) However, no significant differences were observed between the CD20-cells of T2D patients and healthy controls (**Figure 4B and 4D**).

**Figure 4.**
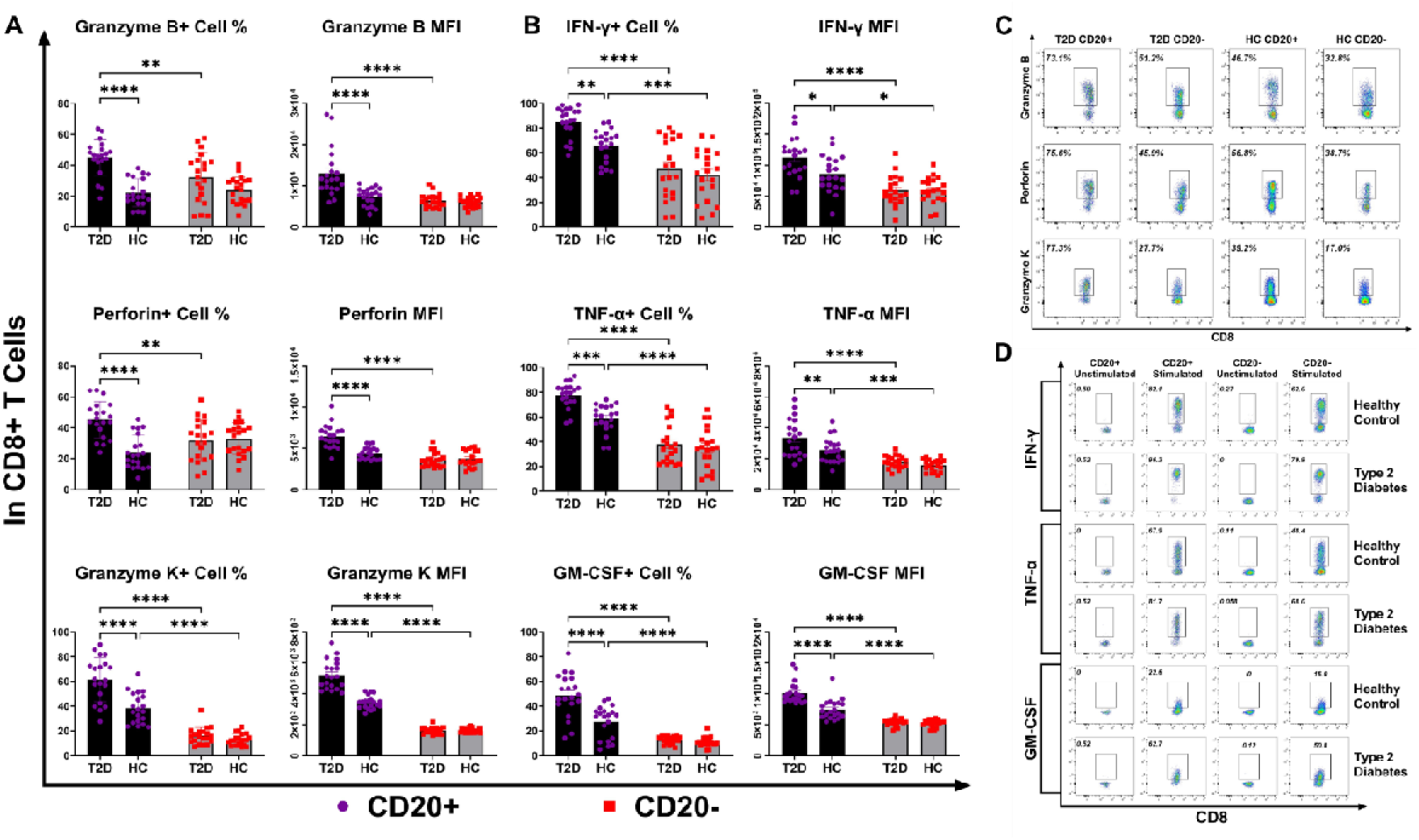
Figure presents the comparison of cell frequencies, and MFI values of **A and C)** Granzyme B, Perforin, Granzyme K; **B and D)** IFN-γ, TNF-α and GM-CSF among CD8+CD20+ and CD8+CD20-subsets of T2D patients and healthy controls, respectively. Statistical analyses were performed using Two-Way ANOVA test with Bonferroni correction. *\*p<0.05, **p<0.01, ***p<0.001, ****p<0.*0001.

### CD20+ Cytotoxic T cells of T2D Patients Have Elevated Proliferative Capacities

To investigate the proliferative capacity of CD20+ and CD20-CTLs in the context of T2D, we analyzed the percentage of proliferating CD8+CD20+ T cells in Healthy Controls and T2D patients.

Firstly, CD20+ CTLs of T2D patients exhibited a markedly enhanced proliferative capacity compared to healthy controls (p<0.0001). In both healthy controls and T2D patients, the percentage of proliferating CD20+ CTLs was significantly higher than that of CD20-counterparts (p<0.001 and p<0.0001 respectively). In contrast, no significant difference in the percentage of proliferation was observed between CD20-CTLs from healthy controls and T2D patients (**Figure 5A and 5B**).

**Figure 5.**
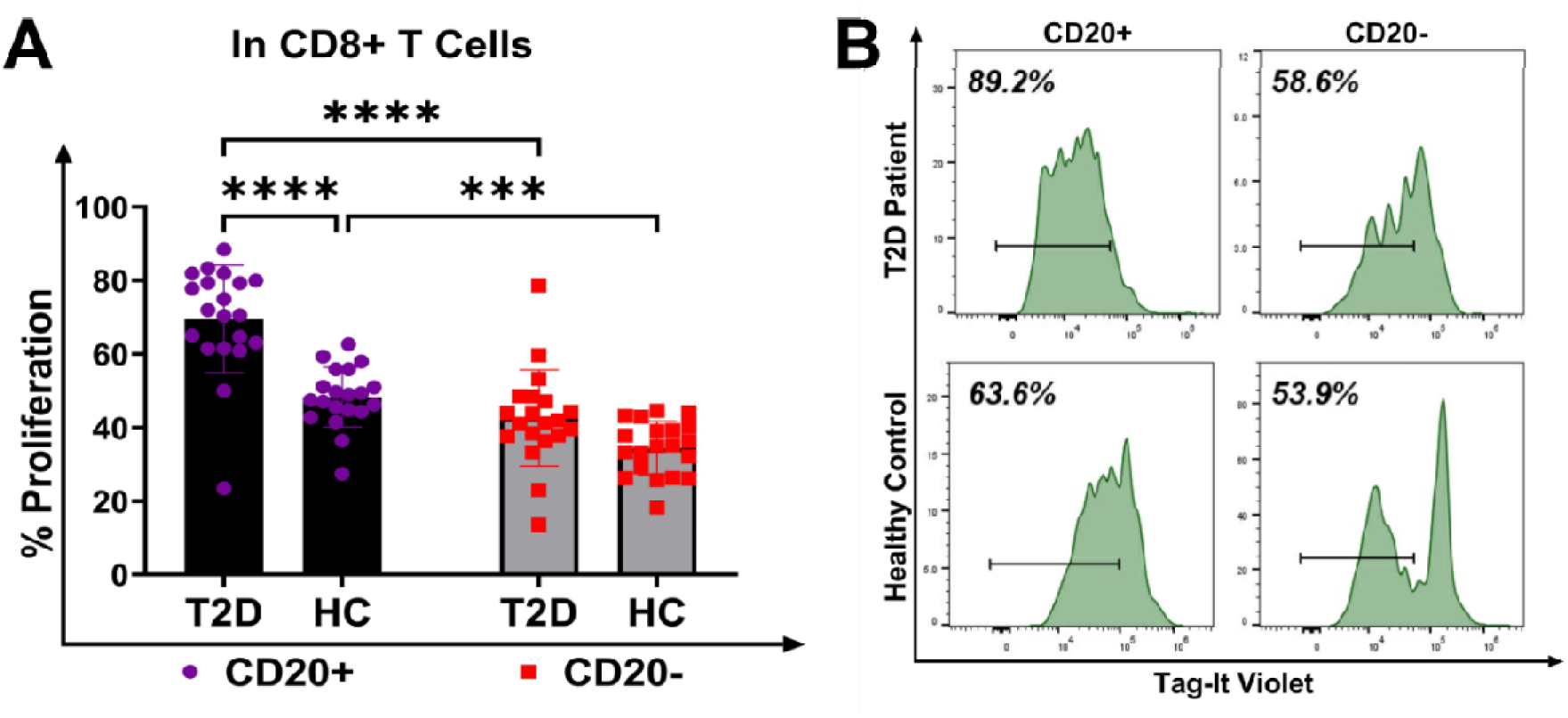
CD20+ CD8+ T cells exhibit higher proliferation in Type 2 Diabetes compared to Healthy Controls. Statistical analysis was performed by Two-Way ANOVA with Bonferroni correction. **A)** Figure shows the percentage of proliferating CD20+ CTLs from Healthy Controls and T2D Patients. **B)** Representative flow cytometric proliferation analysis. *\*p<0.05, **p<0.01, ***p<0.001, ****p<0.*0001.

These results suggest that i) CD8+CD20+ cells are a highly proliferative subset of CTLs in both healthy individuals and those with T2D and ii) the heightened proliferation of CD20+ CTLs in T2D patients suggests a potential role for this specific T cell subset in the pathogenesis or progression of the disease.

### Glucose Induces Dose-Dependent Alterations in the CD8+CD20+ Cell Population and CD20 Expression on CTLs in Healthy Control PBMCs

When we exposed PBMCs from healthy controls to increasing glucose concentrations, we observed a significant increase in both the frequency of CD8LCD20□ cytotoxic T cells and the CD20 expression within these cells across the 100–400 mg/dL glucose range. However, when the glucose concentration was further elevated to 450 mg/dL, both the frequency and MFI values showed a significant decrease. These findings indicate that hyperglycemia exerts a direct effect on CD20□ cytotoxic T cells (**Figure 6A and 6B**).

**Figure 6.**
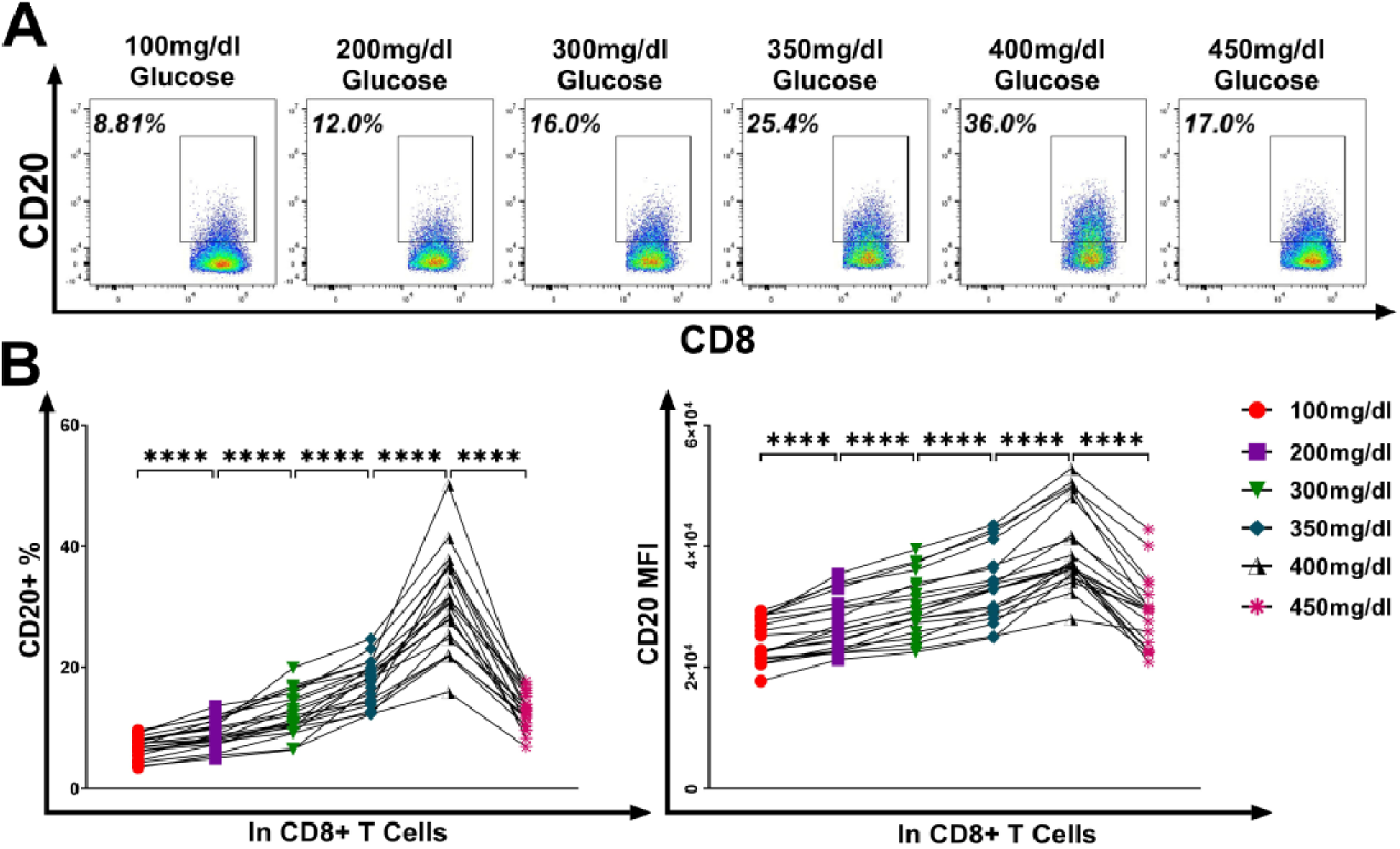
Figure illustrates the alterations on the CD20+ cytotoxic T cell populations and CD20 expression from healthy controls in response to increasing glucose concentrations. Statistical analysis was performed by RM One-Way ANOVA with Bonferroni correction. **A)** A representative graph is provided, showing the changes in the CD8+CD20+ frequencies upon exposure to increasing glucose concentrations. **B)** A dose-dependent elevation was found in both the frequencies of CD20+ cytotoxic T cells and CD20 expression within a glucose range of 100-400 mg/dL. In 450 mg/dL of glucose however, both CD8+CD20+ cell population and CD20 MFI demonstrate a significant decline. *\*p<0.05, **p<0.01, ***p<0.001, ****p<0.*0001.

#### Glucose Availability Enhances Proliferation and Effector Function of CD20□ Cytotoxic T Lymphocytes in a Dose-Dependent Manner

To investigate the influence of glucose availability on CD20+ and CD20-CTL activity, we assessed the proliferation and effector functions of CD8+CD20+ and CD8+CD20-cells of healthy controls cultured under a range of glucose concentrations (100 mg/dL, 200 mg/dL, 300 mg/dL, 350 mg/dL, 400 mg/dL, and 450 mg/dL).

CD8+CD20+ cell proliferation was markedly influenced by glucose concentration. As depicted in the proliferation assay (**Figure 7A, 7B**), a progressive increase in the percentage of proliferating CD20+ CTLs was observed with rising glucose levels, starting from the lowest concentration of 100 mg/dL. Proliferation peaked at a glucose concentration of 400 mg/dL. However, a further increase in glucose to 450 mg/dL resulted in a significant reduction in proliferation. Nevertheless, we found that proliferation in CD20-negative CTLs remained unaffected by increments in glucose concentration.

**Figure 7.**
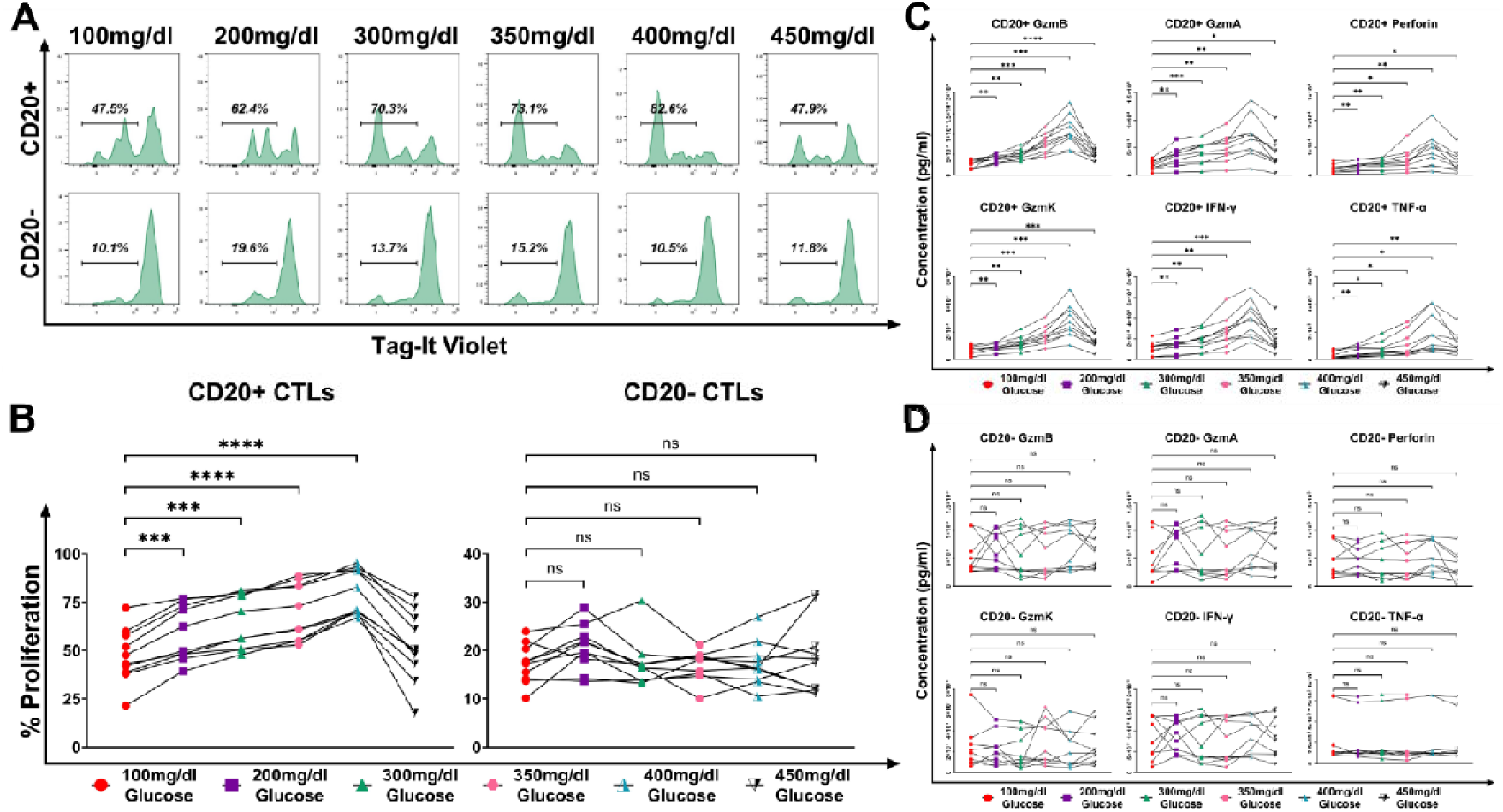
Figure illustrates the proliferative capacity of CD20+ and CD20-cytotoxic T cells from healthy controls in response to increasing glucose concentrations. Statistical analysis was performed by RM One-Way ANOVA with Bonferroni correction. **A)** Representative graph of the proliferative capacities of CD20+ vs. CD20-CTLs in increasing glucose concentrations. **B)** Specifically, CD20+ CTLs exhibit a dose-dependent increase in proliferation as glucose concentrations rise from 100 mg/dL to 400 mg/dL. However, their proliferative capacity demonstrates a decline at a glucose concentration of 450 mg/dL. In contrast, elevation in glucose concentration did not impact the proliferation rates of CD20-negative cytotoxic T lymphocytes. **C)** A dose-dependent increase was noted in the concentrations of Granzyme B, Granzyme A, Granzyme K, Perforin, IFN-γ, and TNF-α produced by CD20+ cytotoxic T cells within a glucose range of 100-400 mg/dL, although this synthesis was suppressed in culture medium containing 450 mg/dL of glucose. **D)** In CD20-CTLs, however, no changes were observed in the concentrations of the aforementioned molecules in response to increasing glucose concentrations. *\*p<0.05, **p<0.01, ***p<0.001, ****p<0.*0001.

Furthermore, in our analyses of the supernatants from the same samples, we determined that the concentrations of the cytolytic molecules Granzyme B (p<0.05), Granzyme A (p<0.05), Granzyme K (p<0.05), and Perforin (p<0.05), as well as the proinflammatory cytokines IFN-γ (p<0.05) and TNF-α (p<0.05) produced by CD20+ CTLs, increased in correlation with rising glucose concentrations from 100 mg/dL to 400 mg/dL. However, at a glucose concentration of 450 mg/dL, we found that the production of aforementioned molecules was diminished (**Figure 7C)**. Conversely, we found that the concentrations of these molecules in CD20-CTLs remained independent of glucose levels (**Figure 7D**).

To ascertain if the observed decrease in proliferation and functions of CD20+ CTLs at 450 mg/dL was a result of glucotoxicity, we compared cell viability across all tested glucose concentrations. Our analysis revealed no significant differences in viability, suggesting that the changes were not due to a general cytotoxic effect of high glucose (**Figure S5A and S5B**).

#### MS4A1□ CD8□ T Cells Exhibit a GZMK-Biased Effector Memory Phenotype in T2D

Subclustering of CD8□ T cells identified a distinct GZMK^high^ cluster in which MS4A1 was significantly upregulated (log□ FC = 1.74, adj. p<0.0001) (**Figure 8A and 8B**). Differential gene-expression analysis between MS4A1□ and MS4A1□ CD8□ T cells confirmed enrichment of the GZMK□ effector signature, with GZMK showing the largest fold-increase (log□ FC = 2.12, adj. p<0.0001). Upregulation of granule-trafficking genes RAB27A (log□ FC = 1.45, adj. p<0.001) and LYST (log□ FC = 1.28, adj. p=0.002), together with CST7 (log□ FC = 1.51, adj. p=0.004), indicated enhanced vesicular transport and granzyme regulation. GZMA, the closest homolog of GZMK, was likewise increased (log□ FC = 1.67, adj. p=0.003). Activation-associated transcripts were markedly elevated, including HLA-DRB1 (log□ FC = 2.06), HLA-DPA1, HLA-DPB1, and HLA-DQB1 (all adj. p<0.001) and the invariant chain CD74 (log□ FC = 2.41, adj. p<0.0001). Members of the AP-1 transcription factor family (JUN, JUNB, JUND, FOS, FOSB) were upregulated (log□ FC 1.3–2.0, adj. p<0.01), consistent with recent activation. Surface-activation genes CD69 (log□ FC = 1.78), FAS/CD95 (log□ FC = 1.36), and CD44 (log□ FC = 1.22) further supported an early-to-mid activation phenotype. MS4A1□ CD8□ T cells showed significant enrichment in chemokines and receptors mediating inflammatory recruitment, including CXCR3 (log□ FC = 1.59), CXCR4 (1.42), GPR183/EBI2 (1.25), and CCL5 (2.05). Genes involved in tissue egress and homing, such as S1PR1 (1.31), S1PR4 (1.21), and ITGA4 (1.27), were concurrently elevated. An effector-memory-like phenotype was evident, with increased KLRG1 (1.63) and CD27 (1.44) alongside survival cytokine receptors IL7R/CD127 (1.38) and IL2RB (1.35). The transcription factors EOMES (1.56) and TCF7 (1.49) were also significantly upregulated (adj. p<0.01), while SELL/CD62L was reduced (log□ FC = –1.47, adj. p=0.003), yielding a KLRG1□ EOMES□ TCF7□ IL7R□ SELL□ pattern characteristic of GZMK□ effector memory T cells (**Figure 8C**). Pathway-enrichment analysis (Enrichr) of DEGs revealed over-representation of T-cell receptor signaling (adj. p<0.0001), antigen processing and presentation (adj. p<0.0001), inflammatory response (adj. p<0.0001), and cytokine–cytokine receptor interaction (adj. p<0.0001) pathways, confirming that MS4A1□ CD8□ T cells represent a recently activated, GZMK-biased effector subset (**Figure 8D**).

**Figure 8.**
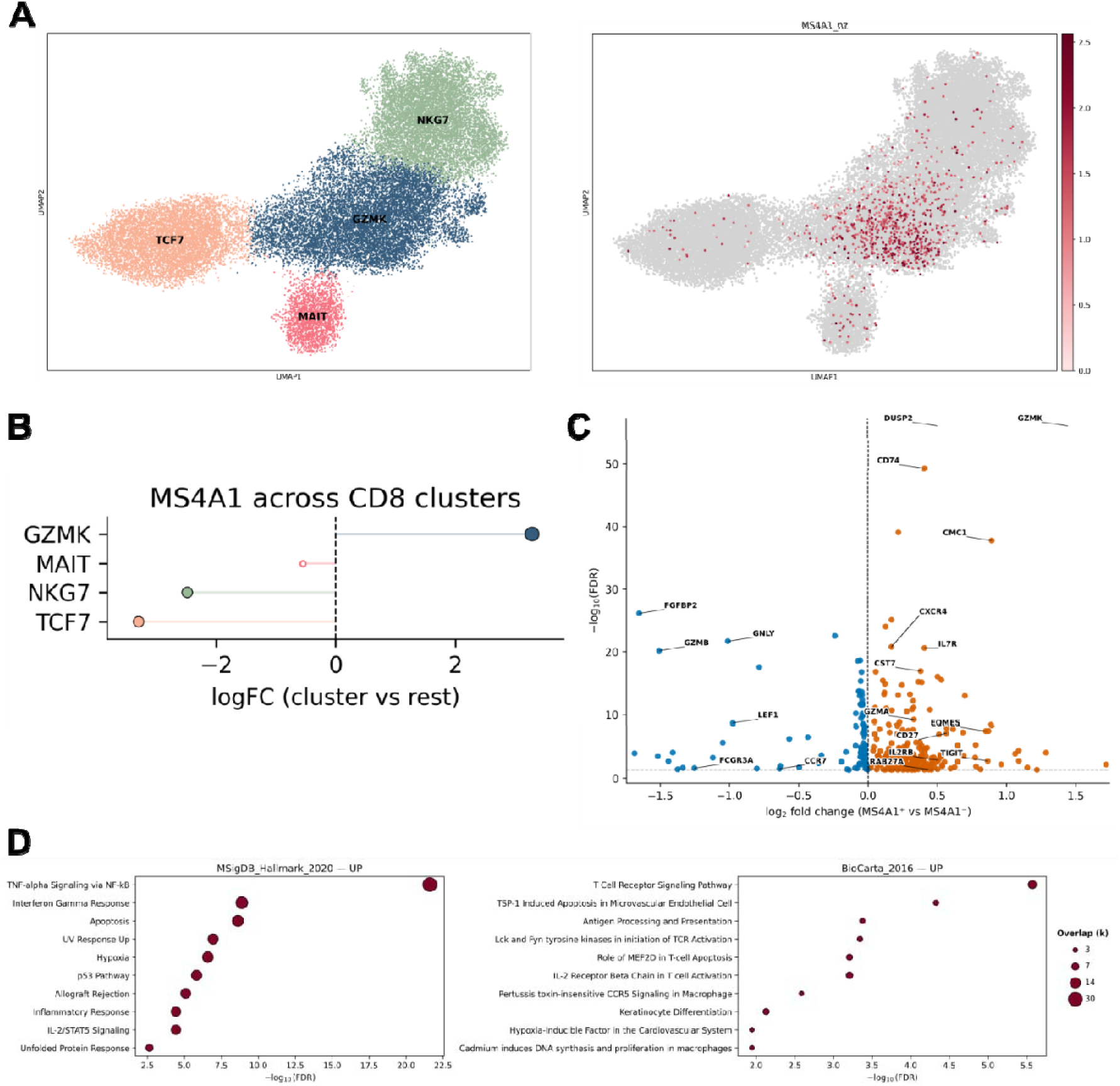
Figure illustrates the DGE of MS4A1+ cytotoxic T cells against MS4A1-counterparts from the scRNA-seq dataset. A) Cytotoxic T cells were subclustered based on their gene expression into TCF7, MAIT, GZMK, and NKG7 clusters. MS4A1+ cells (shown as bordeux dots) were observed to be more densely concentrated in the GZMK cluster. B) Distribution of MS4A1+ cytotoxic T cells within the CD8+ cytotoxic T cell clusters is shown. C) Volcano plot illustrates the genes that are upregulated (shown in orange) and downregulated (shown in blue) in MS4A1+ cytotoxic T cells relative to MS4A1-cells. D) Pathway enrichment analyses of MS4A1L cytotoxic T cells based on the MSigDB Hallmark 2020 and BioCarta 2016 libraries are presented.

#### Effector Memory (EM) CTLs Are Preferentially Expanded and Show the Highest CD20□ Enrichment in T2D

Guided by our single-cell RNA-sequencing (scRNA-seq) analyses, we further characterized CD8□ cytotoxic T-cell (CTL) memory subsets and observed a marked redistribution in individuals with type 2 diabetes (T2D) compared with healthy controls (HC) (**Figure 9A and 9B**). T2D patients displayed significantly higher proportions of effector memory (EM; CCR7□CD45RO□) and terminally differentiated effector memory (EMRA; CCR7□CD45RO□) CTLs (31.32 ± 2.15% and 56.33 ± 2.33%, respectively) than HCs (20.91 ± 1.47% and 39.64 ± 2.73%; p = 0.0017 and p < 0.0001, respectively), while the proportion of naïve CTLs (CCR7□CD45RO□) was significantly lower in T2D patients (16.22 ± 2.07%) than in HCs (34.88 ± 3.02%; p < 0.0001). Central memory CTLs (CM; CCR7□CD45RO□) remained infrequent and comparable between groups (2.14 ± 0.32% vs. 1.09 ± 0.20%; p > 0.9999). Consistent with these quantitative findings, t-SNE projections demonstrated preferential clustering of EM and EMRA CTLs in the T2D group (**Figure 9B**).

**Figure 9.**
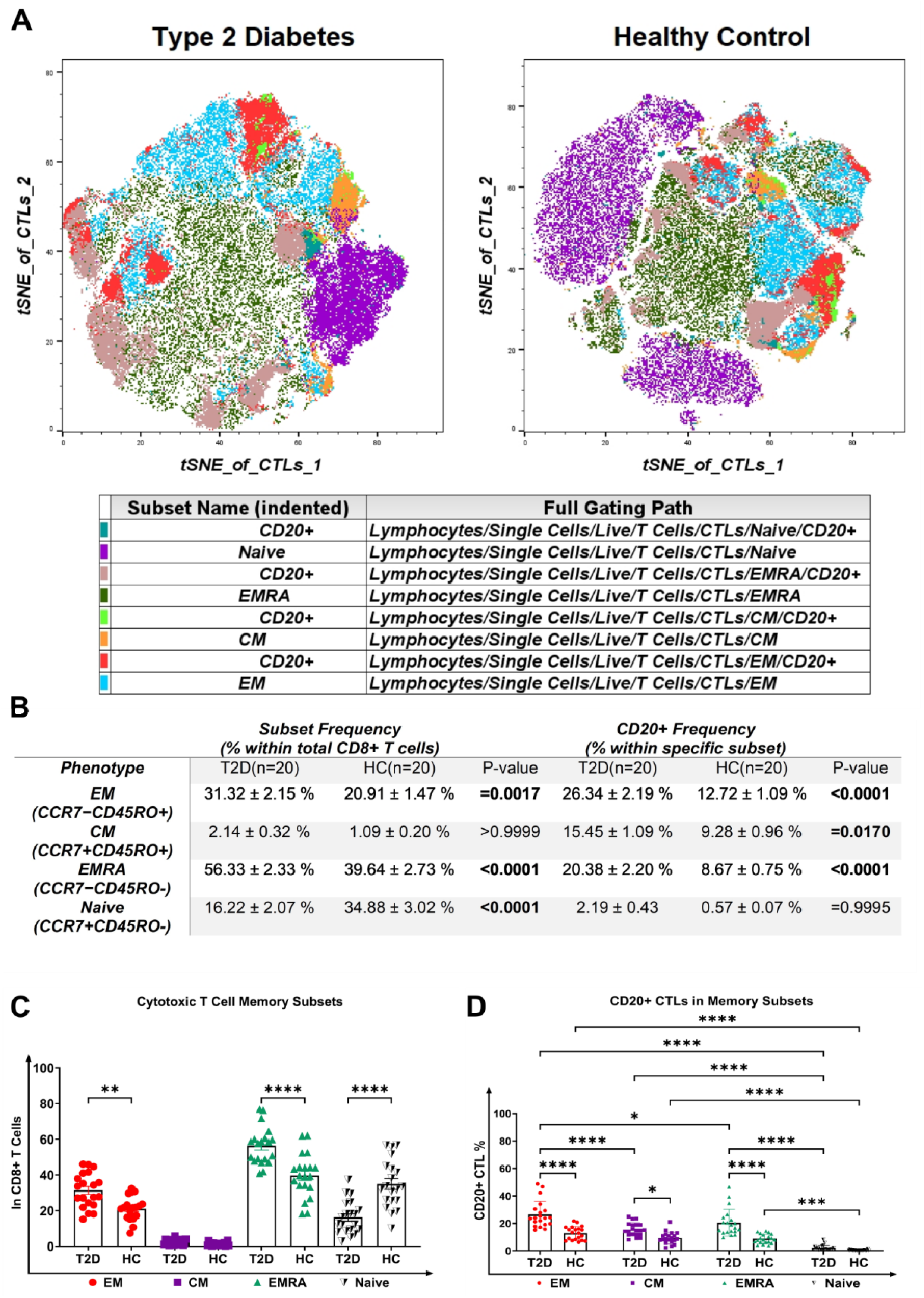
Distribution of CD8□ cytotoxic T-cell (CTL) memory subsets and enrichment of CD20□ CTLs within these subsets in T2D and healthy controls. (A–B) Flow cytometric quantification of CD8L□ CTL memory subsets in peripheral blood from individuals with type 2 diabetes (T2D) and healthy controls (HC). Memory phenotypes were defined as naïve (CCR7□CD45RO□), central memory (CM; CCR7□CD45RO□), effector memory (EM; CCR7□CD45RO□), and terminally differentiated effector memory (EMRA; CCR7□CD45RO□). T2D patients exhibited a significant shift toward differentiated EM and EMRA subsets, with a concomitant reduction in naïve CTLs. Group comparisons were performed using One-Way ANOVA with Bonferroni’s correction. (C–D) Frequency of CD20□ cells within each CD8□ CTL memory subset in T2D and HC. CD20 expression was markedly enriched within EM and EMRA CTLs, with the highest CD20□ frequency observed in the EM subset in T2D. Statistical analysis was performed using Two-Way ANOVA with Bonferroni’s correction to account for both group (T2D vs HC) and memory subset effects. Data are presented as mean ± SEM. *p < 0.05; **p < 0.01; ***p < 0.001; ****p < 0.0001.

We next examined the distribution of CD20□ CTLs within each memory subset and observed a significant enrichment of CD20 expression among EM, CM, and EMRA CTLs in T2D patients compared with HCs (EM: 26.34 ± 2.19% vs. 12.72 ± 1.09%, p < 0.0001; CM: 15.45 ± 1.09% vs. 9.28 ± 0.96%, p = 0.0170; EMRA: 20.38 ± 2.20% vs. 8.67 ± 0.75%, p < 0.0001), whereas CD20□ cells remained rare within the naïve CTL compartment and did not differ between groups (2.19 ± 0.43% vs. 0.57 ± 0.07%; p = 0.9995) (**Figure 9C and 9D**).

Together, these data indicate that T2D is associated with both a shift toward differentiated CTL memory states and a preferential expansion of CD20-expressing CTLs within pro-inflammatory effector pools.

#### CD20□ Cells Are Preferentially Enriched in Granzyme K□ EM and EMRA CD8□ CTLs in T2D

We next investigated whether CD20 expression associates with distinct cytotoxic programs within differentiated CD8□ T-cell memory subsets. We quantified the frequency of CD20□ cells among Granzyme K–positive (GzmK□), Granzyme B–positive (GzmB□), double-positive (GzmK□GzmB□), and double-negative (DN) CTLs within the EM and EMRA compartments in individuals with T2D and healthy controls (**Figure 10A and 10B**). Across both memory subsets, GzmK□ CTLs consistently showed the highest CD20□ frequencies, whereas GrB□ CTLs showed the lowest CD20+ population. CD20□ cells also appeared more frequently in GzmK□ CTLs than in GzmK□GzmB□ and DN populations, indicating that CD20 expression preferentially marks GzmK-skewed effector phenotypes. When we analyzed T2D and HC groups separately, we observed the same pattern: GzmK□ EM and EMRA CTLs contained the highest CD20□ frequencies, with the enrichment appearing more pronounced in T2D. These findings demonstrate that CD20 expression preferentially associates with GzmK-biased EM and EMRA CTLs, rather than with GzmB-dominant cytotoxic subsets.

**Figure 10.**
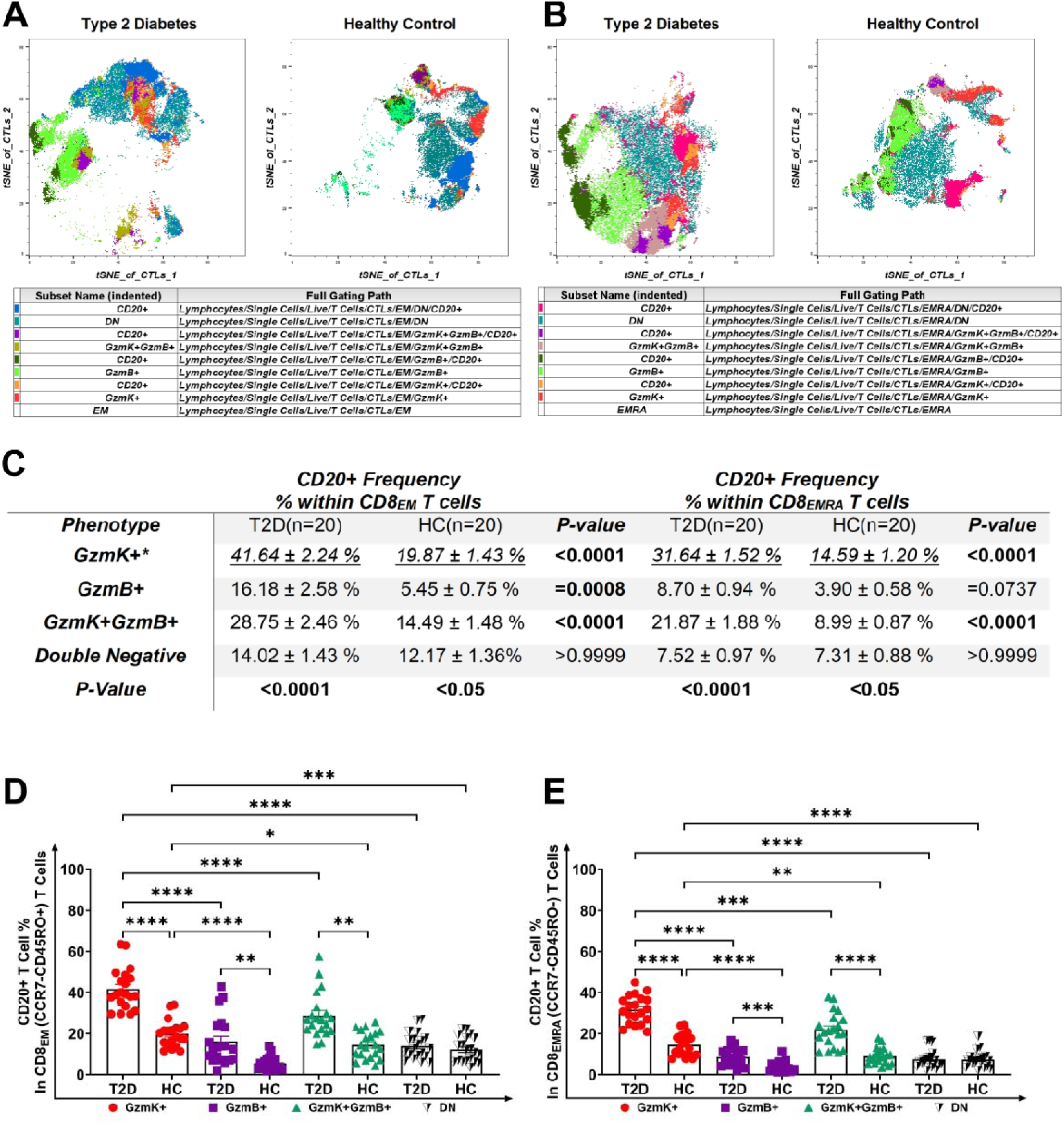
CD20□ enrichment across Granzyme-defined EM and EMRA CD8□ TL subsets in T2D and healthy controls. (A–B) Frequency of CD20□ cells within Granzyme K–positive (GrKL), Granzyme B–positive (GrBL), double-positive (GrKLGrB□), and double-negative (DN) CD8□ cytotoxic T cells (CTLs) in the effector memory (EM) (A) and terminally differentiated effector memory (EMRA) (B) subsets from individuals with type 2 diabetes (T2D) and healthy controls (HC). Across both memory subsets, CD20□ cells were most abundant in GrK□ CTLs, whereas GrB□ CTLs showed the lowest CD20□ frequencies, with intermediate levels in GrKLGrB□ and DN CTLs. The same pattern was evident in both T2D and HC groups, with a more pronounced CD20□ enrichment in GrK□ EM and EMRA CTLs in T2D. Group comparisons within each disease category were performed using One-Way ANOVA with Bonferronis’s Correction. Data are shown as mean ± SEM. *p < 0.05; **p < 0.01; ***p < 0.001; ****p < 0.0001.

## DISCUSSION

Our study provides the first comprehensive functional characterization of CD20+ cytotoxic T lymphocytes in Type 2 diabetes, revealing a distinct immunometabolic axis linking hyperglycemia to adaptive immune dysfunction. In this study, we demonstrated a significant increase in CD20+ T cell populations in the peripheral blood of T2D patients compared to healthy controls. CD20+ T cells, particularly cytotoxic T cells, showed strong correlations with key metabolic and glycemic markers such as HbA1c, fasting blood glucose, LDL, and triglycerides. Functional assays demonstrated increased degranulation and heightened effector molecule production in CD20□ CTLs from T2D patients. These findings highlight the hyperfunctional nature of CD20+ cytotoxic T cells in T2D and suggest their involvement in chronic inflammation and disease progression.

Beyond cytotoxic populations, CD20□ CD4□ T helper cells have also emerged as key immune mediators in chronic inflammatory conditions. These cells exhibit a Th1-skewed phenotype, secrete IFN-γ and GM-CSF, and have been implicated in autoimmune diseases such as multiple sclerosis, Hashimoto’s thyroiditis, and type 3 autoimmune polyendocrine syndrome (12, 14, 15). In this study, CD20□ cytotoxic T cells demonstrated a distinct activation profile in T2D, characterized by higher effector activity compared with both healthy controls and CD20□ CTLs. To our knowledge, circulating CD20□ CTLs have not previously been examined in the context of T2D, and our findings indicate that this subset is selectively expanded and functionally enhanced in metabolic disease. The enhanced proliferative capacity of CD20+ CTLs in T2D patients compared to healthy controls further supports their involvement in disease progression. The positive correlations between CD20+ CTL percentages and metabolic parameters (HbA1c, FBG, LDL, triglycerides) reinforce their clinical relevance. Recent studies have shown that hyperglycemia and dyslipidemia can directly modulate T-cell signaling, cytotoxic granule production, and inflammatory cytokine release (26). These cells may not only contribute to systemic inflammation but also potentially to localized inflammation within metabolic tissues, such as adipose tissue or the liver, thereby directly influencing insulin sensitivity and glucose homeostasis. In a study by Pinho et al., which is relevant to our research, increased percentages of CD20+ helper and cytotoxic T cells were observed in the peripheral blood and orbital adipose tissues of obese patients with insulin resistance. These cells were shown to be polarized towards TH-1 and TC-1 phenotypes and produced higher levels of IFN-γ compared to non-obese patients. Based on these findings, the authors suggested that CD20+ T cells may exhibit hyperfunctionality based dysfunction in type 2 diabetes (16). Unlike previous phenotypic studies, our functional profiling establishes a direct link between CD20□ CTLs and inflammatory capacity in T2D.

Granzyme K is increasingly recognized as a marker of tissue-homing, pro-inflammatory effector T cells rather than classical cytotoxic mediators. A study by Jonsson et al. demonstrated that Granzyme K+ CD8 cytotoxic T cells are highly abundant in the synovial fluid of patients with rheumatoid arthritis and that Granzyme K promotes the synthesis of proinflammatory cytokines and chemokines (27). Similarly, a study by Keiserman et al. showed that Granzyme K induces the secretion of IL-6 and IL-8 from endothelial cells by activating the PAR-2 receptor (28). Studies also have shown that Granzyme K enhances the effects of IFN-γ, a cytokine abundantly secreted by Granzyme K+ cytotoxic T cells (29, 30) Moreover, there are studies in the literature indicating that Granzyme K is involved in diseases characterized by chronic inflammation, such as rheumatoid arthritis, systemic lupus erythematosus, Sjögren’s syndrome, systemic sclerosis, and uveitis (27). In our study, the enrichment of Granzyme K cells within the CD20□ CTL subset—together with their glucose-responsive increase in Granzyme K release—suggests that this population may be particularly sensitive to metabolic stress and capable of contributing to local inflammatory activity in T2D.

In our in vitro glucose exposure experiments, CD20□ CTLs demonstrated a glucose□concentration□dependent biphasic response: from 100 to ∼400 mg/dL, both proliferation and effector functions increased, whereas at 450 mg/dL there was a notable drop in function despite preserved viability. This pattern aligns with emerging concepts of metabolic T cell exhaustion in diabetes, where chronic hyperglycemic exposure drives initial hyperactivation followed by dysfunction. Evidence from literature suggest that in type 2 diabetes, CD8□PD-1□ T cells exhibit impaired metabolic□immune axis coupling, with reduced multifunctionality and cytokine secretion under hyperglycemic conditions despite not necessarily showing increased cell death, consistent with exhaustion rather than cytotoxic loss (31), (32), (33). Thus, our findings extend prior observations by demonstrating that functional decline in CTLs occurs only once glucose surpasses a threshold, aligning with the transitional loss of effector capacity seen in T2D literature, potentially mediated by upregulation of inhibitory receptors, and altered metabolic signaling. Prior studies also have shown that glucose activates NFAT and mTOR-dependent cytokine pathways in effector T cells (34, 35). Our data suggests that hyperglycemia promotes CD20□ CTL activation, but sustained extreme levels may drive exhaustion or dysfunction.

A key insight from our single-cell analyses was the close association between MS4A1 expression and a GZMK-enriched CD8□ T-cell identity. The predominance of GZMK, together with its known co-expression partner GZMA,(36) suggests that CD20□ CTLs in T2D adopt an effector-memory–like program previously linked to tissue-directed inflammatory activity. Crucially, this GZMK-biased identity is not merely a transcriptional signature but is supported by evidence of functional cytotoxic potential. We observed the upregulation of RAB27A and LYST, genes essential for lytic granule trafficking (37–39), as well as CST7, which modulates granzyme activation (40). This suggests that the MS4A1□ population expanded in T2D patients is equipped with the necessary machinery to execute granzyme-mediated functions. Consistent with their GZMK-enriched identity, MS4A1□ CD8 T cells displayed transcriptional features of recent activation, including increased expression of MHC class II genes and AP-1 family members (28), (40–42). The co-expression of CD69, FAS, and CD44 further supported an activated effector state (43–45). In parallel, the upregulation of CXCR3, CXCR4, GPR183, and CCL5 suggests a migratory, tissue-homing potential characteristic of GZMK□ effector subsets (40), (46–49). Moreover, enrichment of KLRG1, CD27, IL7R, and the transcription factor EOMES indicates an effector-memory–like program with potential for persistence within inflammatory microenvironments (50–56). Taken together, these molecular signatures place MS4A1□ CD8 T cells within the broader framework of GZMK-driven inflammatory effector-memory T cells, aligning with their expansion in T2D.

The identification of hyperfunctional CD20+ CTLs in T2D opens novel therapeutic avenues for precision immunomodulation in metabolic disease. While anti-CD20 therapies like rituximab have shown efficacy in preserving β-cell function in Type 1 diabetes and treating severe insulin resistance syndromes, their application in T2D remains largely unexplored. Its application in T2D is not well-established; however, there are instances where rituximab has been used to address severe insulin resistance. For example, a patient with type B insulin resistance was successfully treated using rituximab in combination with cyclophosphamide and prednisone, with significant improvements in glycemic control (57, 58). These observations raise the possibility that modulation of CD20□ immune populations may influence glucose metabolism and insulin sensitivity, particularly in autoimmune-associated contexts. However, whether such therapeutic strategies would benefit the broader T2D population remains unknown. Further studies are needed to determine whether selective targeting of CD20□ T cells—distinct from conventional B-cell depletion—could offer a novel immunotherapeutic approach in T2D.

Several important limitations warrant consideration when interpreting our findings. First, our restriction to treatment-naïve T2D patients, while eliminating medication confounding, resulted in a relatively modest sample size (n=25 per group) that may not capture the full heterogeneity of T2D presentations across diverse populations and disease stages. Second, the cross-sectional design precludes causal inference; longitudinal studies tracking CD20+ CTL dynamics from prediabetes through established T2D are essential to determine whether these cells drive disease progression or emerge as a consequence of metabolic dysfunction. Third, our exclusive analysis of peripheral blood may not fully represent tissue-resident CD20+ CTL populations in metabolically active organs (adipose tissue, liver, pancreas) where local inflammation drives insulin resistance and β-cell dysfunction. Fourth, while our in vitro glucose exposure experiments provide mechanistic insights, they cannot recapitulate the complex metabolic milieu of T2D, where multiple factors including insulin resistance, dyslipidemia, adipokines, and inflammatory mediators interact synergistically.

Finally, the lack of follow-up data on patient outcomes limits our ability to assess whether CD20+ CTL levels correlate with disease progression, treatment response, or the development of diabetic complications over time. The reversibility of CD20+ CTL expansion and dysfunction with metabolic improvement (through lifestyle intervention, bariatric surgery, or pharmacotherapy) could establish these cells as biomarkers for treatment response and guide personalized therapy.

## CONCLUSIONS

In summary, this study identifies CD20□ cytotoxic T lymphocytes as a metabolically responsive, hyperfunctional T cell subset that may contribute to the chronic low-grade inflammation characteristic of type 2 diabetes. The increased frequency; enhanced cytotoxic and proinflammatory activity; and positive correlations of CD20□ CTLs with HbA1c, fasting glucose, LDL, and triglycerides highlight their potential involvement in disease pathogenesis. The observed glucose-dependent biphasic response—marked by progressive activation up to 400 mg/dL and a subsequent functional decline at 450 mg/dL despite preserved viability—suggests that excessive metabolic stimulation may trigger an exhaustion-like state. These findings imply that moderate hyperglycemia promotes CD20□ CTL activation and effector function, whereas extreme glucose levels may lead to metabolic dysregulation and immune dysfunction. The detailed transcriptional profile of MS4A1□ CD8 T cells demonstrates a comprehensive overlap with the established signature of GZMK□ T cells. This includes not only the expression of granzymes but also the full phenotypic program of activation, inflammatory migration, and an EOMES-driven effector memory state. When viewed in the context of our primary finding — that CD8□CD20□ T cells are expanded in T2D and correlate with metabolic dysfunction — this analysis provides a crucial mechanistic link. The data strongly suggest that the “CD20□ CTL” population identified in our T2D cohort is, in fact, this GZMK-biased, highly activated, and persistent memory population. Therefore, MS4A1 (CD20) expression emerges as more than a novel marker; it identifies a key pathogenic cell population, suggesting that CD20□ CTLs are a novel and highly relevant player in the chronic inflammation underlying the immunopathogenesis of Type 2 Diabetes.

Collectively, our results, even as preliminary, reveal a novel link between metabolic stress and CD20+ CTL activation in type 2 diabetes and underscore the need for future studies to determine whether targeted modulation of CD20□ CTLs can mitigate inflammation and improve metabolic homeostasis.

## LIST of SUPPLEMENTARY MATERIALS

Figure S1 to S56

**Tables S1**

## ACKNOWLEDGEMENTS

We would like to thank the patients and healthy volunteers who participated in this research project. The authors gratefully acknowledge the use of the services and facilities of the Koç University Research Center for Translational Medicine (KUTTAM), funded by the Presidency of Turkey, Presidency of Strategy and Budget. During the preparation of this work the authors used Gemini 3.0 for English language editing of this article. After using this tool, the authors reviewed and edited the content as needed and took full responsibility for the content of the publication.

## AUTHOR CONTRIBUTIONS

Conceptualization: ÖA

Design of experiments: ÖA

Investigation: ÖA, SSK, SU, AB, AV

Formal analysis: ÖA, AV, SSK, SU, AB

Resources: AB, DY, OD, SÇD, SÖ, NA, BM, ZYY, HS

Funding acquisition: ÖA

Visualization: AV, ÖA, SU

Supervision: AV

Writing – original draft: ÖA, AV, AB

Writing – review & editing: AB, OD, DY

## DATA and MATERIALS AVILABILITY

All data are available in the main text or the supplementary materials.

