## Supplementary Figures for "Glucose-Responsive CD20+ Cytotoxic T Cells: A Novel Pro-inflammatory Mediator in the Immunopathogenesis of Type 2 Diabetes"

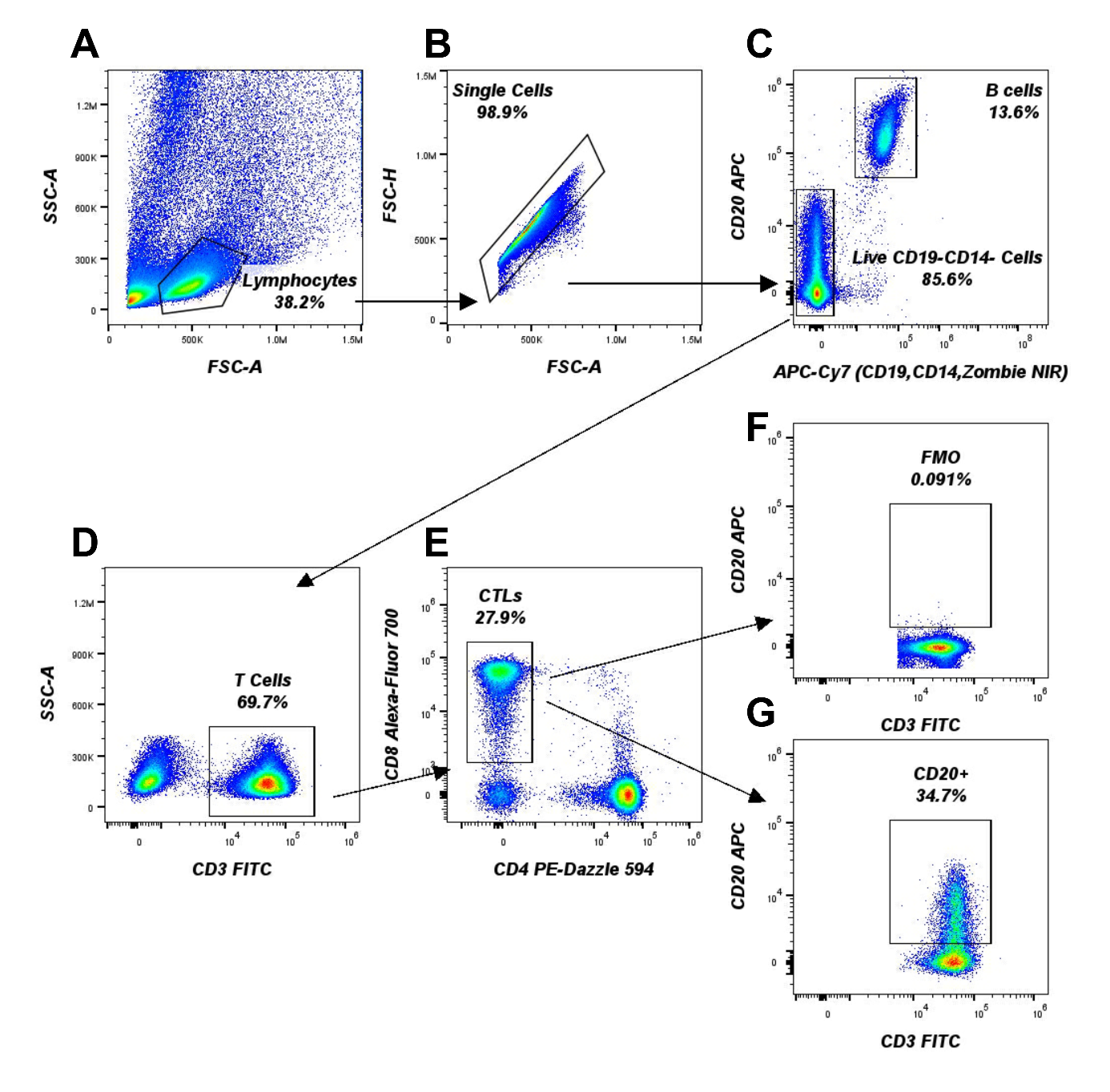

**Figure S1.** Gating strategy for CD20⁺ cytotoxic T cells. **(A)** Lymphocytes were gated based on the FSC-A × SSC-A plot, followed by **(B)** selection of single cells by excluding doublets using the FSC-A × FSC-H plot. **(C)** CD20⁺ lymphocytes were gated on the APC-Cy7 (CD19, CD14, Zombie-NIR) × CD20 APC plot by excluding B cells, monocytes, and dead cells. **(D)** T cells were gated on the CD3 FITC × SSC-A plot, and **(E)** CD8⁺CD4⁻ cells were identified as CTLs based on the CD4 PE-Dazzle 594 × CD8 AF700 plot. **(F)** CD20 FMO. **(G)** CD20⁺ cytotoxic T cells.

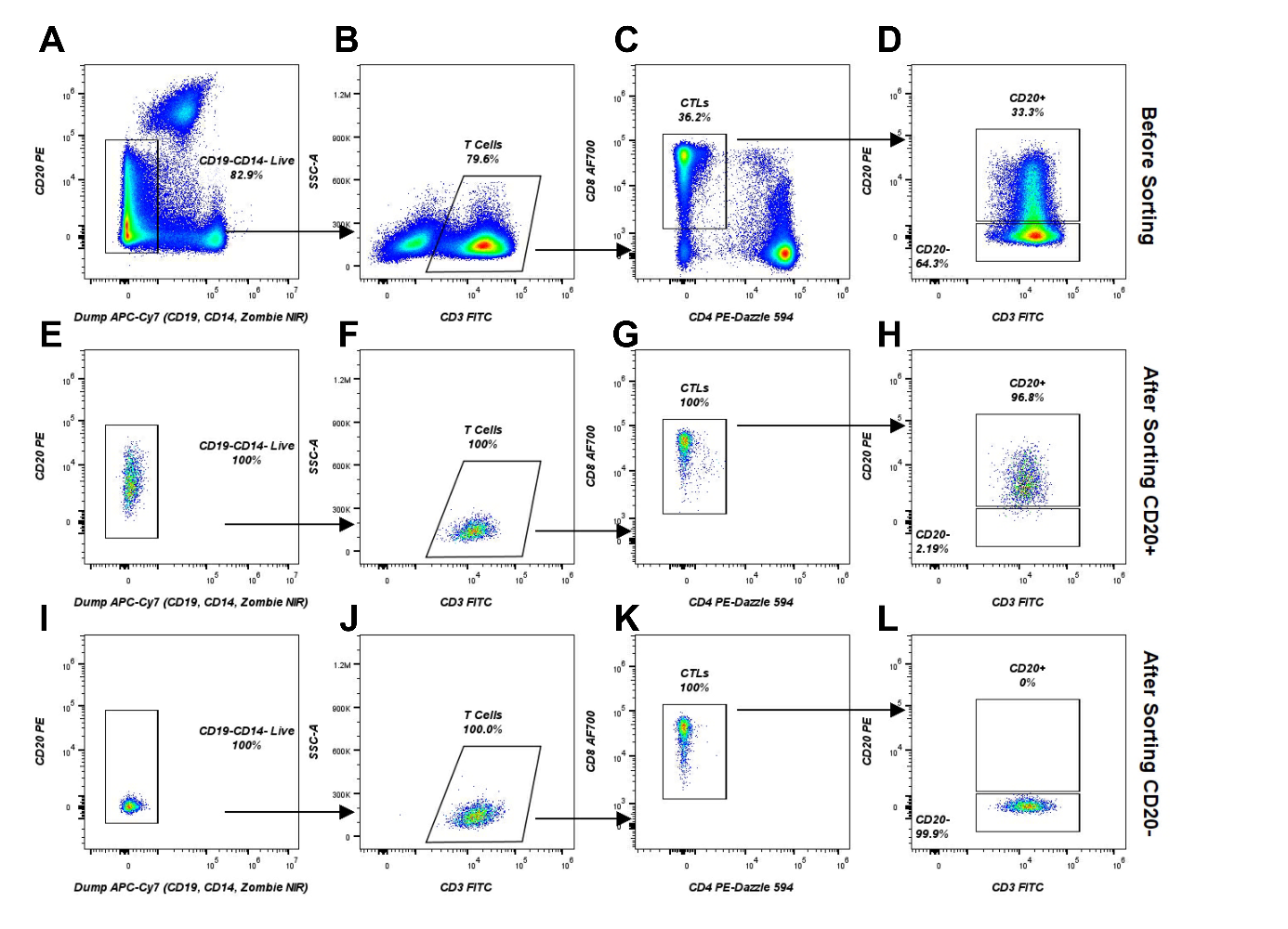

**Figure S2. Isolation of CD20+ and CD20- T cell subsets by fluorescence-activated cell sorting (FACS).** Gating strategy and post-sort purity analysis are shown. The pre-sort sample was first gated on live, CD19- and CD14- Cells **(A)**, followed by gating for CD3+ T cells **(B)** and CD8+ CTLs **(C)**. The starting population contained 23.3% CD20+ and 64.2% CD20- T cells **(D)**. **(E-H)** After sorting for the CD20+ population, purity analysis showed that the collected fraction was 96.8% CD20+. (I-L) The fraction sorted for CD20- cells was confirmed to be 99.9% pure.

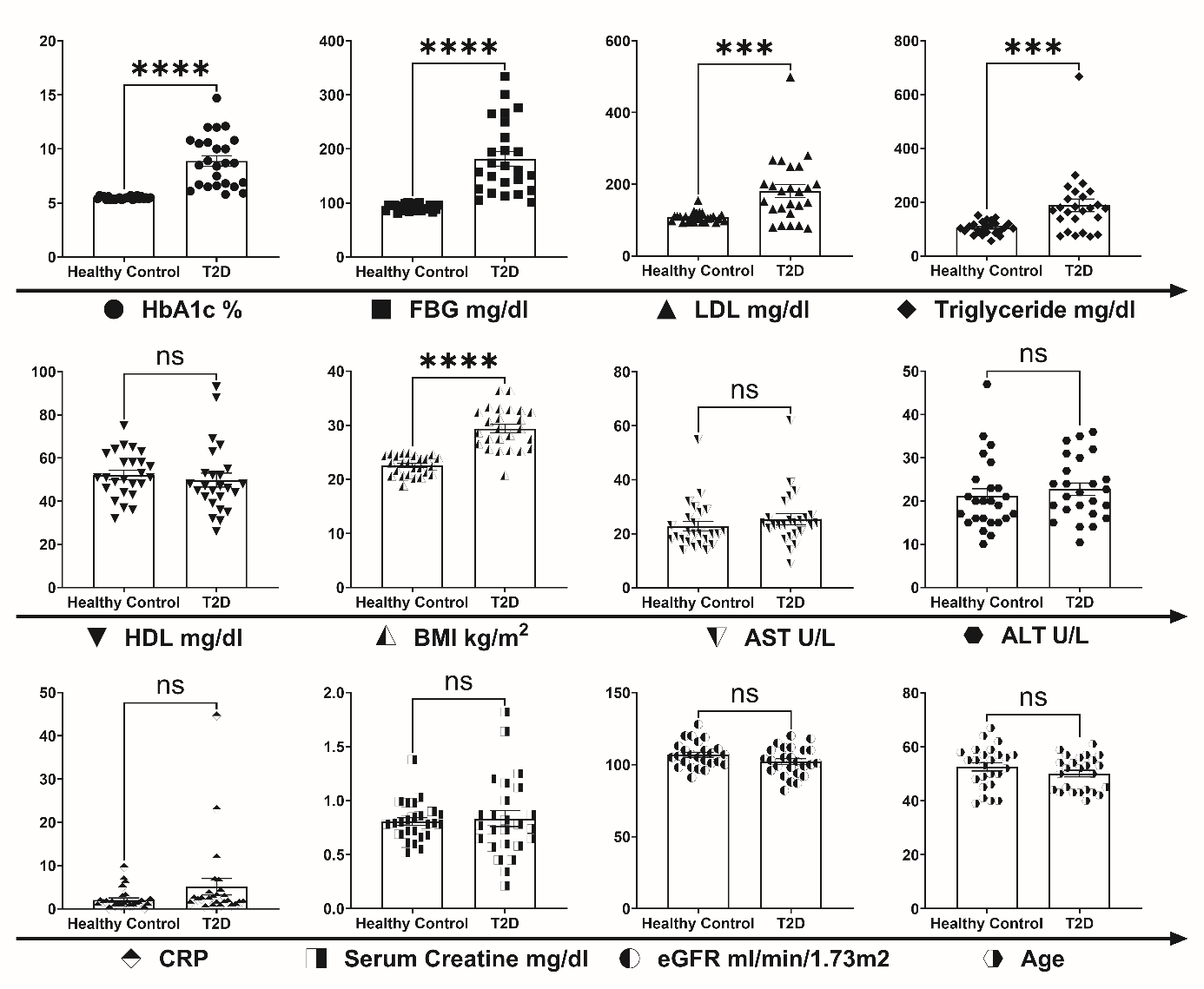

**Figure S3.** Figure presents the statistical analysis comparing the clinical parameters of patients with T2D and the healthy controls enrolled in the study. Mann-Whitney U test was performed for statistical analyses .**p<0.05, **p<0.01, ***p<0.001, ****p<0.0001***.**

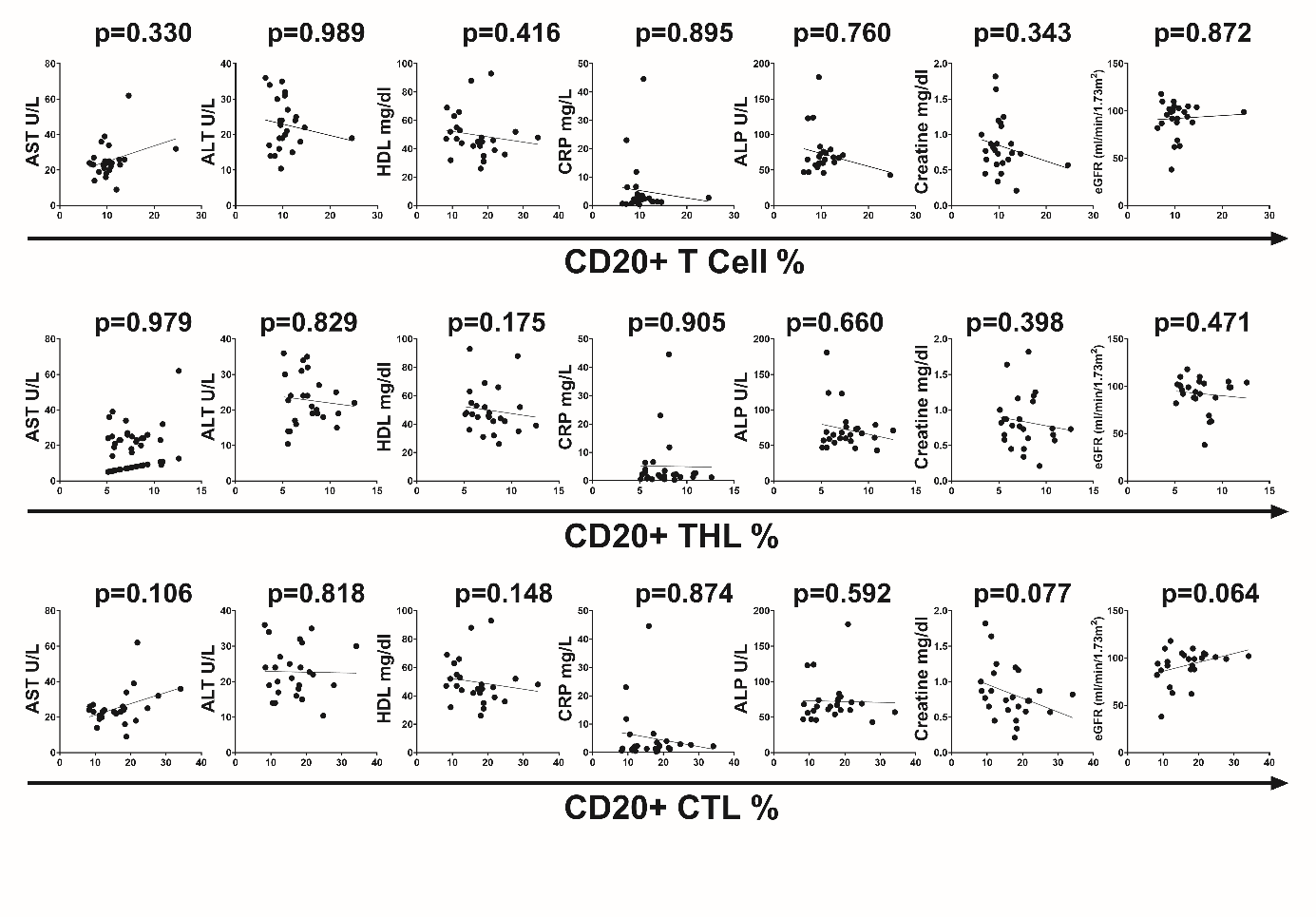

**Figure S4.** Figure presents the correlation analyses between the clinical parameters of T2D patients and the percentages of CD20+ T cells. Non-parametric Spearman test was utilized for the analysis. *ns* *p>0.05*

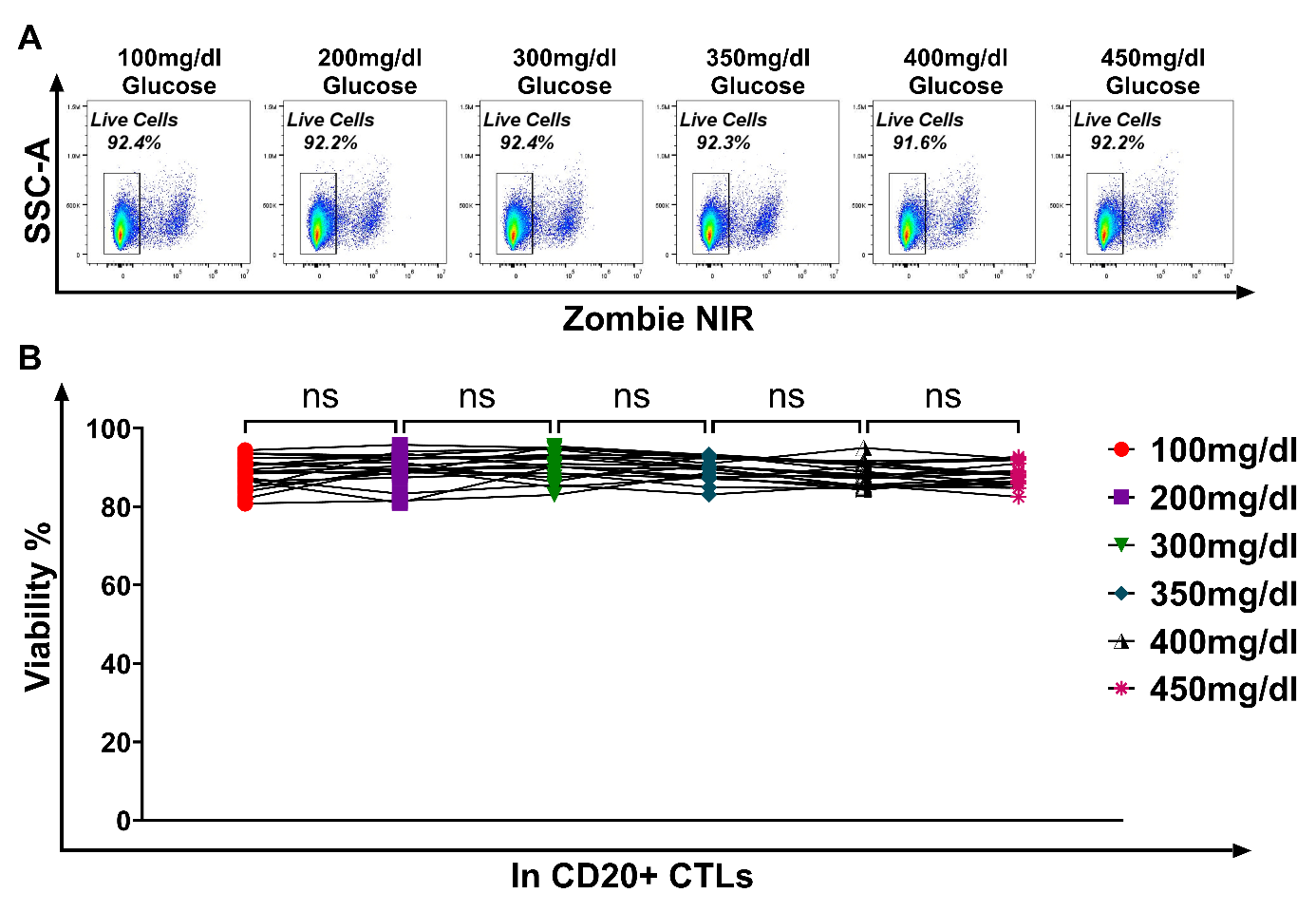

**Figure S5.** Assessment of glucotoxicity in CD8+CD20+ cells exposed to hyperglycemia. The viability of CD8+CD20+ cells from healthy controls was compared across increasing glucose concentrations to determine if hyperglycemia induces glucotoxicity. Statistical analyses were performed using a Repeated Measures One-Way ANOVA test. **A)** A representative plot of cell viability. **B)** No significant change was observed in the viability of CD8+CD20+ cells with increasing glucose concentrations. *ns* *p>0.05*

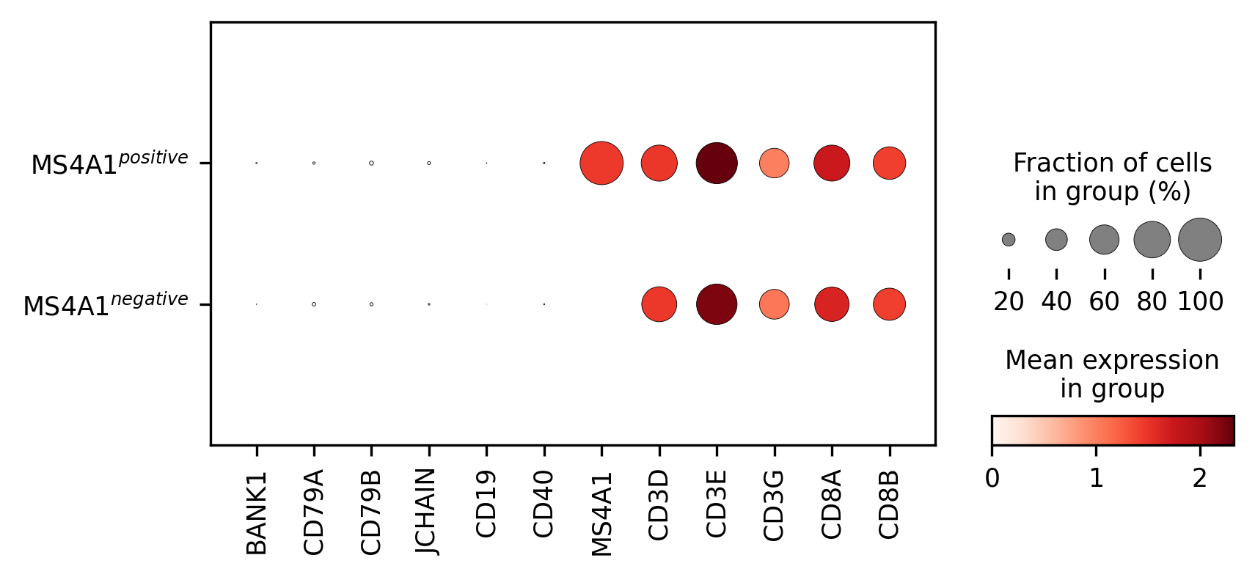

**Figure S6.** Figure illustrates the identification of MS4A1+ cytotoxic T cells from the scRNA-seq dataset. After B cells and all B cell-associated genes were excluded from PBMC samples, MS4A1+ and MS4A1- cytotoxic T cells were identified.
